# CHD7 interacts with the nucleosome acidic patch for its efficient activity via its N-terminal region

**DOI:** 10.1101/2020.11.19.389429

**Authors:** Eunhye Lee, Chanshin Kang, Pasi Purhonen, Hans Hebert, Karim Bouazoune, Sungchul Hohng, Ji-Joon Song

## Abstract

Chromodomain-Helicase DNA binding protein 7 (CHD7) is an ATP dependent chromatin remodeler involved in maintaining open chromatin structure. Mutations of CHD7 gene causes multiple developmental disorders, notably CHARGE syndrome. However, there is not much known about the molecular mechanism by which CHD7 remodels nucleosomes. Here, we performed integrative biophysical analysis on CHD7 chromatin remodeler using crosslinking-mass spectrometry (XL-MS), cryo-Electron Microscopy (cryo-EM) and single-molecule Förster Resonance Energy Transfer (smFRET). We uncover that N-terminal to the Chromodomain (N-CRD) interacts with nucleosome. Importantly, this region is required for efficient ATPase stimulation and nucleosome remodeling activity of CHD7. The cryo-EM analysis on the N-CRD_Chromodomain bound to nucleosome reveals that the N-CRD interacts with the acidic patch of nucleosome. Furthermore, smFRET analysis shows the mutations in the N-CRD result in slow or highly-fluctuating remodeling activity. Collectively, our results uncover the functional importance of a previously unidentified N-terminal region in CHD7 and implicate that the multiple domains in chromatin remodelers are involved in regulating their activities.

## Introduction

Eukaryotic genome is hierarchically organized into high-order structure called chromatin. Chromatin is composed of arrays of nucleosomes, which consist of 146 base pairs of DNA wrapped around a histone octamer (1). This chromatin organization restricts the access of protein factors to DNA. Therefore, cellular processes such as transcription, DNA replication or DNA repair require dynamic reorganization of the chromatin structure (2,3). To this end, cells devised several ways to overcome this challenge. Among these, histone modification and ATP-dependent chromatin remodeling processes play a major role in reorganizing the chromatin structure and these two processes are functionally interconnected (4–6). The N-terminal tails of histone are modified with several moieties such as acetyl-, phosphor-, ubiquityl-, ADP-ribosyl and methyl-groups, and these modifications function to recruit proteins having reader domains recognizing specific histone modifications (7). For example, Chromodomains and PHD domains are responsible for recognizing methylated lysines of histones (8,9). ATP-dependent chromatin remodelers are responsible for reorganizing chromatin structure by sliding nucleosome, exchanging histones and altering nucleosome structure (10). These remodelers often form large multi-subunit complexes and can be classified into four main families based on their ATPase subunit: mating type SWItch/Sucrose Non-Fermenting (SWI/SNF), Imitation SWItch (ISWI), INOsitol requiring 80/SWi2-Snf2-Related 1 (INO80/SWR1) and Chromodomain Helicase DNA binding (CHD). Among these, the remodelers belonging to CHD family are unique in that they do not form multi-subunit complexes and a ATPase subunit alone is fully functional (10,11). CHD family member can be further categorized into three subfamilies based on their domains (12).

CHD7 is a member of the CHD-subfamily III and contains a double Chromodomain, an ATPase, SANT and BRK domains. The CHD7 chromodomains have been reported to bind methylated histone residues, including histone H3 Lysine 4 (H3K4). Consistent with these observations, genome-wide analyses show that CHD7 occupies enhancer and promoter regions, where methylated H3K4 is enriched (13). The functional importance of the CHD7 chromodomains is further supported by genetic studies linking chromodomain mutations to Kallmann syndrome, Idiopathic Hypogonadotropic Hypogonadism (IHH), and Coloboma of the eye, Heart defects, Atresia of the choanae, Retardation of growth, Genital and Ear abnormalities (CHARGE) syndrome (14–16). In fact, mutations across the entire CHD7 gene are the main cause of CHARGE syndrome, and 30% of the affected new-born die before the age of 5 (17,18). Moreover, CHD7 is frequently mutated in Autism Spectrum Disorders (19). Given that the CHD7 ATPase domain is essential for nucleosome remodeling, disease mutations which fall into this domain are often believed to inactivate the protein. Nevertheless, due to the lack of detailed functional studies on the role of the other CHD7 protein domains, the impact of most patient mutations on CHD7 functions is unknown.

Recent cryo-EM structures of yeast Chd1 (subfamily I) and human CHD4 (subfamily II) provide important hints regarding the molecular mechanism by which CHD family proteins remodel nucleosomes (20,21). In the structures, ATPase domains occupy at Super Helical Location (SHL) +2 position in nucleosome while their Chromodomains are located at SHL+1 and interact with nucleosomal DNA (20,21). However, despite its importance in human disease and development, little has been characterized about CHD7 remodeling activity and nucleosome binding.

Here, we analyze the interaction between full-length CHD7 and nucleosome by cross-linking mass spectrometry (XL-MS) and cryo-electron microscopy (cryo-EM). We find that the N-terminal region to the Chromodomain (N-CRD) is involved in recognizing the acidic patch of the nucleosome. Furthermore, we characterized the contribution of the N-CRD to the CHD7 activity by using single-molecule Föster Resonance Energy Transfer (smFRET) combined with Restriction Enzyme Accessibility (REA) assay, showing that the N-CRD is required for efficient remodeling activity of CHD7 and functions as a remodeling throttle by interacting with the acidic patch. Thus, our work uncovers a new regulatory region in CHD7 and provide a molecular basis for interpreting the impact of disease mutations occurring in proteins of the CHD subfamily III.

## Results

### The CHD7 N-terminal region to Chromodomain (N-CRD) interacts with nucleosome

CHD7 is composed of several domains including double Chromodomain, ATPase, SANT and BRK domains and some of these domains in other remodelers have been involved in interacting with nucleosome (22–24). However, it is not clear if other regions of CHD7 besides the domains already known are involved in recognizing nucleosome and regulating its activity. Therefore, to examine the interaction network between CHD7 and nucleosome, we performed cross-linking mass spectrometry (XL-MS) analysis using disuccinimidyl suberate (DSS). Full-length recombinant CHD7 and nucleosome with 40 bp linker DNA were crosslinked in the presence of 1 mM MgCl_2_ and 1 mM ADP.BeFx. The homogeneity of the cross-linked complexes was analyzed by native PAGE (Supplementary Fig. 1). The XL-MS analysis reveals that the crosslinks with histones are clustered in the region in N-terminal to the Chromodomain (N-CRD) (Fig.1A and Supplementary Table 1), suggesting that the N-CRD of CHD7 is located proximal to nucleosome in the complex. Therefore, we next examined whether N-CRD may directly bind nucleosomes by Electrophoresis Mobility Shift Assay (EMSA) (Fig. 1B, 1C and Supplmentary Fig. 2). The N-CRD_Chromodomain (CD) or the N-CRD alone are sufficient to bind to nucleosomes but not Chromodomain alone, suggesting that the N-CRD directly binds to the nucleosome.

**Figure 1.**
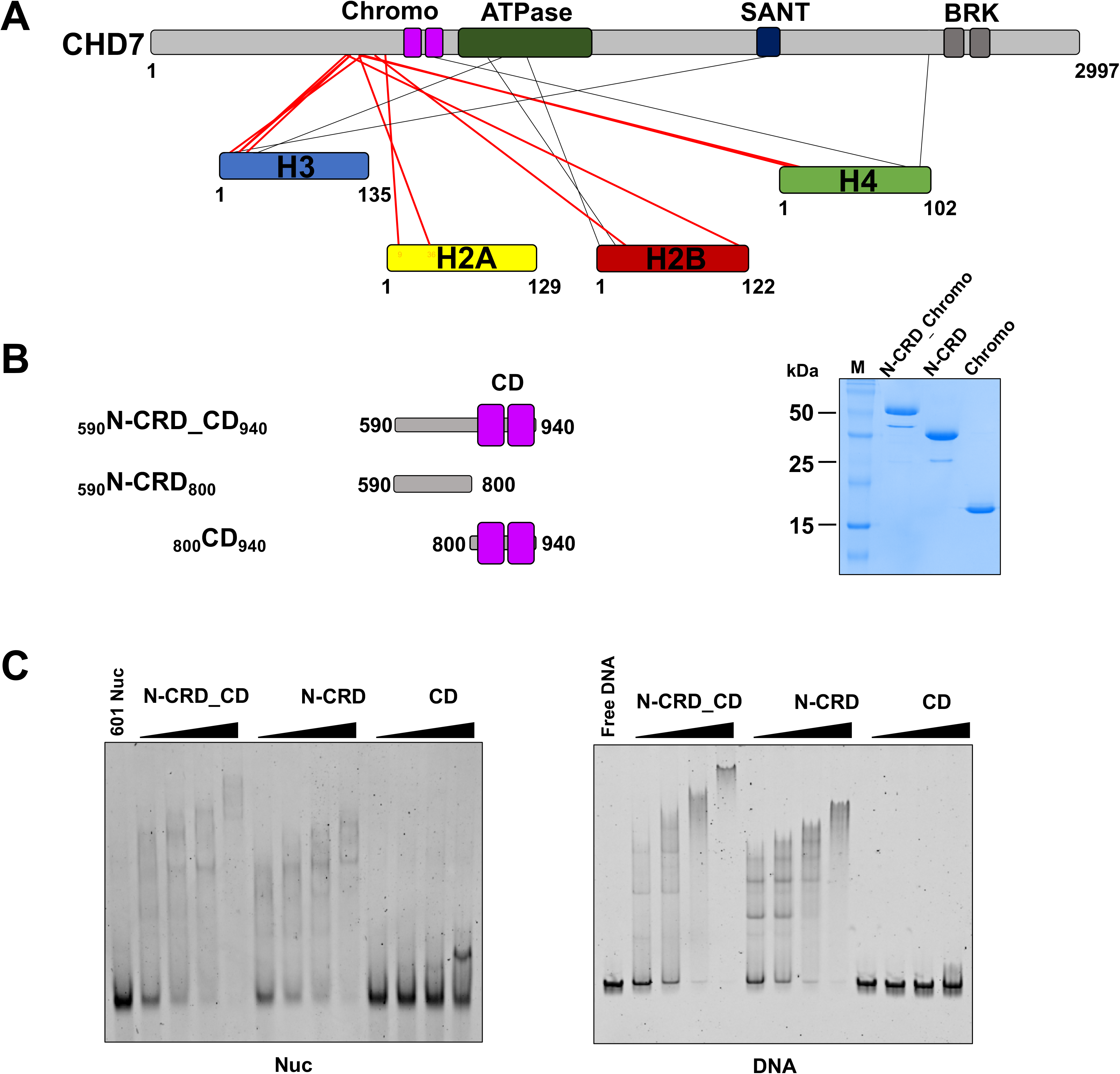
N-terminal to Chromodomain (N-CRD) of CHD7 interacts with nucleosome. (A) Crosslinking-mass spectrometry analysis of full-length CHD7 bound to nucleosome. Thick red lines show the crosslinks between N-CRD and histones. (B) A scheme of CHD7 constructs (_590_N-CRD_CD_940_, _590_N-CRD_800_, _800_CD_940_). CD stands for Chromodomain. The qualities of the proteins were analyzed by SDS-PAGE stained with coomassie blue (right panel). (C) EMSA assay with _590_N-CRD_CD_940_, _590_N-CRD_800_ and _800_CD_940_ with nucleosome (Nuc) (left panel) and 187 bp free DNA (DNA) (right panel). Nucleosome (2.5 μM) or free DNA (1 μM) were incubated with increasing amounts of the proteins. The bindings were visualized by a native PAGE with Red safe staining.

### The N-CRD is required for efficient remodeling activity of CHD7

As our XL-MS and EMSA analysis suggest that the N-CRD is involved in interacting with nucleosome, we examined the role of the N-CRD in CHD7 activities. We generated a deletion construct of full-length CHD7 where N-CRD is deleted (CHD7_ΔN-CRD_) (Fig. 2A). The CHD7_ΔN-CRD_ protein behaved as wild-type CHD7 (CHD7_WT_) during purification, and the limited proteolysis analysis on CHD7_WT_ and CHD7_ΔN-CRD_ shows the similar cleavage pattern, suggesting that the deletion does not disrupt the global structure of CHD7 (Supplementary Fig. 3). It was previously shown that the ATPase activity of CHD7 is stimulated by substrates (25). Therefore, we first examined if the N-CRD deletion affects the substrate-mediated stimulation of ATPase activity (Fig. 2B). In the absence of substrates, the basal ATPase activity of CHD7_ΔN-CRD_ is comparable to that of CHD7_WT_, further indicating that the deletion does not affect the integrity of CHD7. In the presence of DNA, the ATPase activities of CHD7_WT_ and CHD7_ΔN-CRD_ were equally stimulated. However, while CHD7_WT_ ATPase activity is greatly stimulated by nucleosome substrates, the deletion of the N-CRD significantly dampens the nucleosome-mediated stimulation of ATPase activity, emphasizing that the N-CRD is likely involved in nucleosome interactions and critical for the substrate-mediated stimulation of the ATPase activity. Second, we examined the contribution of the N-CRD to the remodeling activity of CHD7 using Restriction Enzyme Accessibility (REA) assay (Fig. 2C). This assay monitores the ability of remodeling factors to expose a restriction site by sliding the histone octamer away from the restriction site. Hence, we measured the percentage of cleavage by the MfeI restriction enzyme, as a result of nucleosome remodeling by CHD7. Quantification of the REA assays shows that the deletion of the N-CRD results in substantial decrease of the CHD7 remodeling activity. These data demonstrate that the N-CRD is required for efficient remodeling activity of CHD7 as well as for propor nucleosome stimulated ATPase activity.

**Figure 2.**
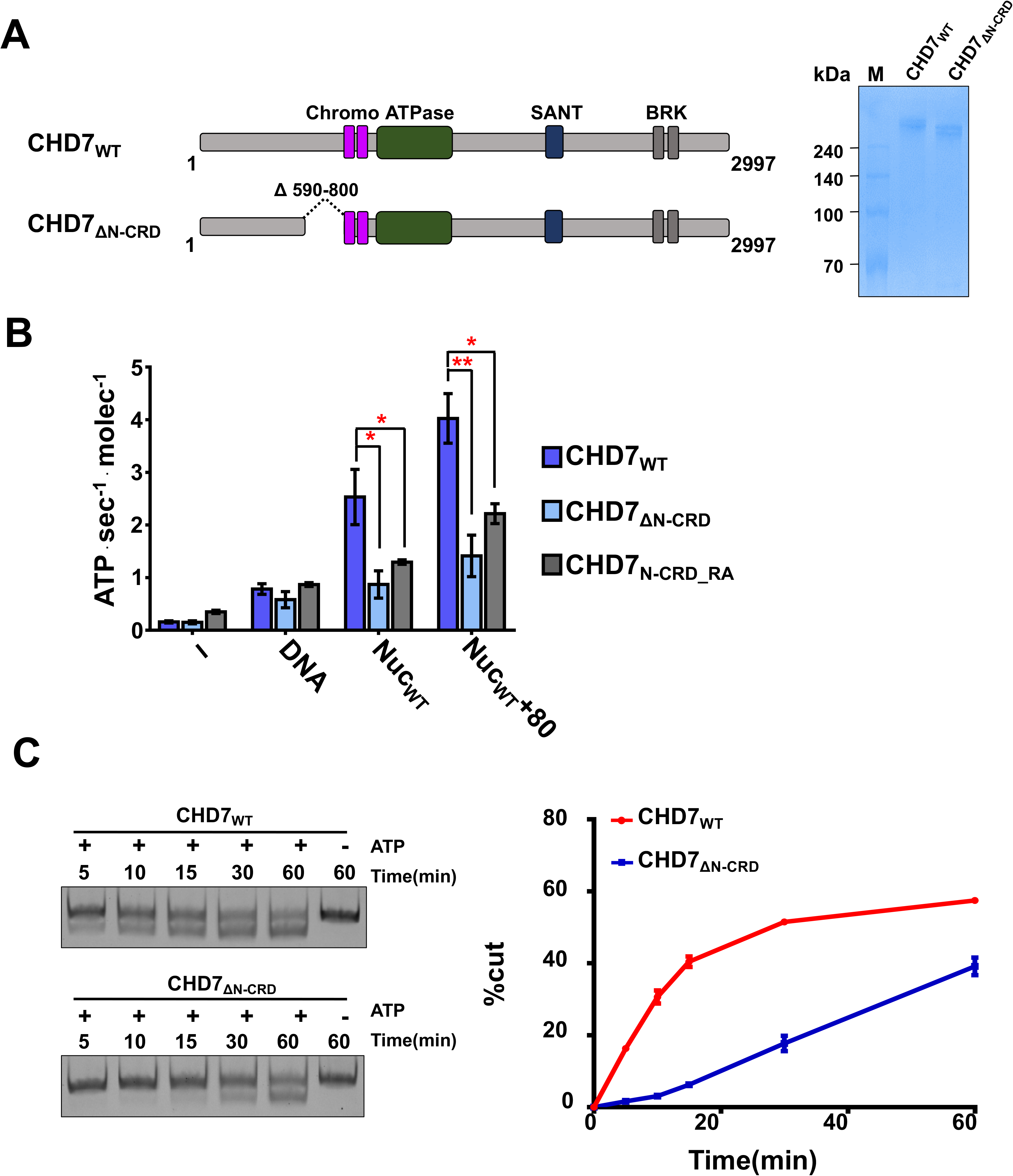
The N-CRD of CHD7 is required for efficient remodeling activity. (A) A schematic representation of wild-type CHD7 (CHD7_WT_), N-CRD deletion mutant (CHD7_ΔN-CRD_) and a Arginine to Alanine point mutant in N-CRD (CHD7_N-CRD_RA_). The qualities of the proteins were analyzed by SDS-PAGE stained with coomassie blue. (B) Substrate mediated stimulation of CHD7 ATPase activity. The ATPase activities of full-length CHD7_WT_ or CHD7_ΔN-CRD_ were measured in the presence of buffer, free DNA, nucleosome core (Nuc_WT_) or nucleosome with 80 bp linker DNA (Nuc_WT_ +80). P-value was calculated with the student-t test from three independent experiments (n=3) (* P< 0.05, ** P<0.0001) (C) Representative gels showing the CHD7 remodeling activity of wild-type CHD7_WT_ and CHD7_ΔN-CRD_ measured by REA assay. Nucleosomes with +80 bp linker DNA were incubated with 50 nM of CHD7_WT_ or CHD7_ΔN-CRD_, and quenched at 5, 10, 15, 30 and 60 mins. The cut DNAs were extracted and analyzed by native PAGE with Red safe staining. The graph shows the quantification of the percentage of cut DNA from three independent experiments (n=3) with standard errors.

### The Cryo-EM structure of the N-CRD_Chromodomain bound to nucleosome reveals its interaction with the acidic patch

As our data show that the N-CRD plays critical roles in the CHD7 activity, we examined if there is any conserved region in the N-CRD by aligning sequences among CHD7s from several species. Among the species compared, fly CHD7 orthologue, Kismet shows very diverged sequences from the other species. However, there is a highly conserved stretch of positively charged residues within the N-CRD, suggesting its functional importance (Fig. 3A).

**Figure 3.**
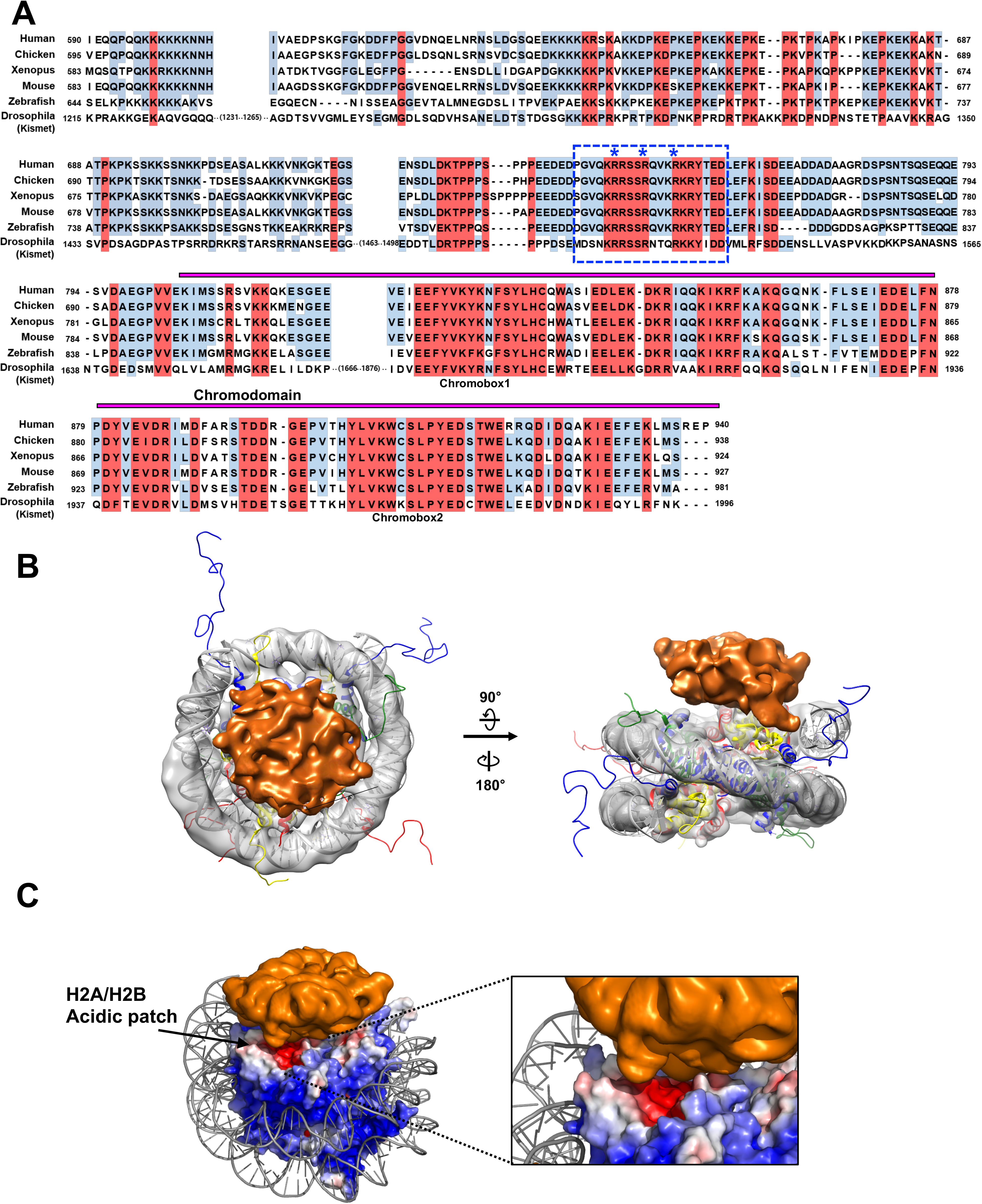
The N-CRD of CHD7 contacts with the acidic patch of nucleosome. (A) A sequence alignment of N-CRD_Chromodomain among CHD7 orthologues. Chromdomain is indicated with a magenta bar above the sequence. The highly conserved arginine stretch is boxed with a blue dotted line. Arginines mutated to alanines are indicated with asterisks. (B) The cryo-EM map of N-CRD_Chromodomain bound to nucleosome. The cryo-EM map for nucleosome is shown in gray and the crystal structure of nucleosome (1KX5) was docked in the map. The extra-density for N-CRD_Chromodomain is shown in orange. The map was drawn at a contour 0.0057 for nucleosome and 0.0021 for N-CRD_Chromodomain respectively. (C) An electrostatic surface representation of nucleosome showing the acidic patch contacting with the cryo-EM density from N-CRD_Chromodomain.

Although XL-MS roughly mapped the interaction between N-CRD and nucleosome, to better understand the interaction between nucleosome and N-CRD, we determined the cryo-EM structure of the N-CRD_Chromodomain bound to nucleosome core (Fig. 3B, 3C, Supplementary Fig. 4 and Supplementary Table 2). Purified N-CRD_Chromodomain bound to 601 nucleosome core was cross-linked with 0.1% glutaraldhyde and further separated by sucrose gradient ultra-centrifugation. The complex was vitrified and a total of 9,369 micrographs were collected using Titan Krios 300KeV with a Gatan K2 Summit direct detector at SciLife Lab, Stockholm. A total of 1,859,258 particles were initially picked and further subjected to multiple rounds of 2D class averages and 3D classifications using Relion 3.0 (26). From a final of 17,743 particles, we obtained a cryo-EM map of N-CRD_Chromodomain-nucleosome complex at 7.25 Å resolution estimated by FSC criterion of 0.143.

Unfortunately, we were not able to obtain a high-resolution cryo-EM structure. However, the cryo-EM map clearly resolved the secondary elements of nucleosome and reveals the extra-density corresponding to N-CRD_CD. The Chromodomain of CHD1 in the cryo-EM structure shows that it positions at SHL+1 and interacts with the ATPase lobe, which is far away from the side surface of nucleosome (21). Therefore, it is likely that the extra density observed in our cryo-EM corresponds to the N-CRD of CHD7. Interestingly, the extra density covers the acidic patch of nucleosome, suggesting the interaction between nucleosome and N-CRD via the acidic patch (Fig. 3C). To further validate that N-CRD interacts with nucleosome, we deteremined the cryo-EM structure of N-CRD without chromodomain bound to nucleosome having 40 bp linker DNA (Supplementary Fig. 5 and Supplementary Fig. 6). In this low resolution cryo-EM structure, N-CRD is located near the acidic patch, which is the exact same location as seen in the N-CRD_CD nucleosome core structure. These two cryo-EM structures strongly indicate that N-CRD is involved in recognizing the acidic patch.

Therefore, we examined if the acidic patch affects CHD7 activity. We mutated acidic residues in histone H2A (E61, E64, D90, E92) to alanines (A) and assembled acidic patch mutant (APM) nucleosome (Nuc_APM_) (Fig. 4A and 4B). We first examined if Nuc_APM_ still stimulates CHD7 ATPase activity. Consistent with the data showing that N-CRD deletion dampens the nucleosome-mediated stimulation and N-CRD interacts with the acidic patch, the level of the activation by Nuc_APM_ substantially decreases compared with NucWT regardless of the presence of a linker DNA (Fig. 4C). We then performed REA assay, showing that CHD7 has almost no activity with Nuc_APM_. (Fig. 4D).

**Figure 4.**
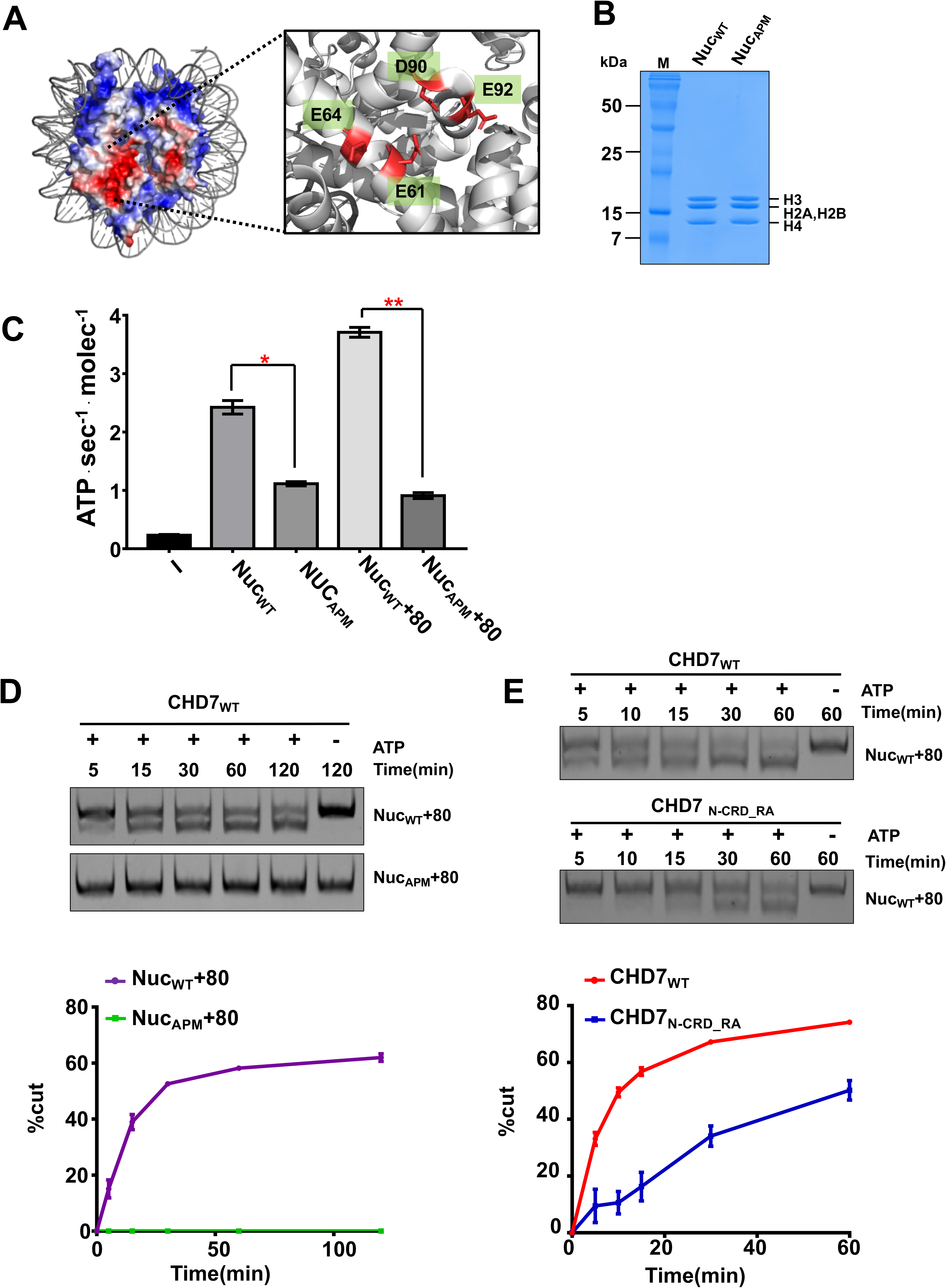
The acidic patch is an important for CHD7 activity. (A) An electrostatic surface representation of the octamer showing the acidic patch. In this study, the E61, E64, D90 and E92 residues were mutated to alanines. (B) Nucleosomes used in the REA assay were analyzed by SDS-PAGE gel followed by coomassie blue staining. (C) The stimulation of CHD7 ATPase activity by Nuc_WT_, Nuc_APM_, Nuc_WT_+80 and Nuc_APM_+80. The error bars were calculated from three independent experiments (n=3). P-value was calculated with the student-t test (*P< 0.0005, ** P<0.0001) (D) The remodeling activity was measured by REA assay. 20 nM of Nuc_WT_+80 and Nuc_APM_+80 were each incubated with 50 nM of CHD7 and quenched at 5 min, 10 min, 30 min, 60 min and 120 min. Reactions were analyzed by native PAGE with Red safe staining. The line graph (left panel) shows quantification of the percentage of cut DNA from three independent experiments with standard error (n=3). (E) Representative the gels showing CHD7_N-CRD_RA_ remodeling activity measured by REA assay. 20 nM of Nuc_WT_+80 were incubated with 50 nM of CHD7 and quenched at 5 min, 10 min, 15 min, 30 min and 60 min. Reactions were analyzed by native PAGE with Red safe staining. The line graph (left panel) shows the quantification of the percentage of cut DNA from three independent experiments with standard error (n=3).

The current low resolution structure did not allow to map the exact region in the N-CRD interacting with the acidic patch. However, as arginine anchor composed of several arginines is shown to interact with the acidic patch of nucleosome in several nucleosome binding proteins (27–35), we examined if there is an arginine anchor motif in N-CRD. Interestingly, highly conserved arginines are clustered in N-CRD suggesting the existance of a possible arginine anchor in N-CRD (Fig. 3A). Therefore, we mutated some of these arginines (R748, R752 and R756) to alanines in full-length CHD7 (CHD7_N-CRD_RA_) (Supplementary Fig. 7). As CHD7_ΔN-CRD_ has substantially lost the ATPase stimulation activity by nucleosome, we examined if it is the case for CHD7_N-CRD_RA_ (Fig. 2B). The ATPase assay shows that CHD7_N-CRD_RA_ behaves as CHD7_ΔN-CRD_ in that the stimulation of ATPase activity by nucleosome is greatly dampened, suggesting that the functional important role of the arginines in N-CRD (Fig. 2B). We then measured the remodeling activity of CHD7_N-CRD_RA_ (Fig. 4E). The REA assay with the arginine mutant CHD7_N-CRD_RA_ shows substantial decrease in the remodeling activity, suggesting that the arginine stretch at N-CRD might function as an arginine anchor interacting with nucleosome.

### Single-molecule FRET analysis shows that the N-CRD plays a critical role for efficient remodeling activity of CHD7

In order to further characterize the contribution of the N-CRD to CHD7 activity, we studied the CHD7 remodeling activity using smFRET. We prepared a nucleosome composed of 601 DNA sequence, 3 bp linker on the exit side, and 78 bp linker on the entry side (Fig. 5A and Supplementary Fig. 8A). Cy5 and Cy3 are labeled at the end of the exit side DNA, and T120C of H2A of the histone octamer, respectively (36) so that high FRET efficiency is obtained before remodeling. The DNA is also labeled with a biotin at the end of the entry side DNA for surface immobilization. After immobilizing nucleosomes on a polymer-coated quartz surface using the streptavidin-biotin interaction, we measured FRET efficiencies of individual nucleosomes. The immobilized nucleosomes exhibited three FRET peaks, each corresponding to the three different labeled species (Supplementary Fig. 8B). Of these three species, we used for data analysis only the high FRET species in which a single Cy3 is attached to a position proximal to the exit side. Therefore, FRET decrease could be used to monitor the translocation of DNA at the exit side. To monitor the remodeling activity of CHD7 on the immobilized nucleosomes, we sequentially injected CHD7 and ATP. When the wild type CHD7 (200 nM) was injected into a chamber, clear Protein Induced Fluorescence Enhancement (PIFE) signals were observed from many (76.5 %) immobilized nucleosomes (Fig. 5B). These PIFE intensity were 20 % greater than an intensity before CHD7 injection (Fig. 5C). After washing out free CHD7 to prevent multiple binding/dissociation events of CHD7, we added ATP to initiate the remodeling. Remodeling events (stepwise FRET decrease) were observed only from the molecules that previously exhibited PIFE (Fig. 5B), indicating that CHD7 binding can be monitored using PIFE occurrence. From 134 molecules that exhibited the stepwise FRET decrease, FRET histograms before the remodeling (P0, Fig. 5B) and after the remodeling (P1, Fig. 5B) were made. The width of P1 was similar to that of P0, suggesting that CHD7 remodels a nucleosome with a well-defined kinetic step size (Fig. 5D). Based on the calibration data (Supplementary Fig. 9), the kinetic remodeling step size of CHD7 was estimated as 5.9 bp (Fig. 5E).

**Figure 5.**
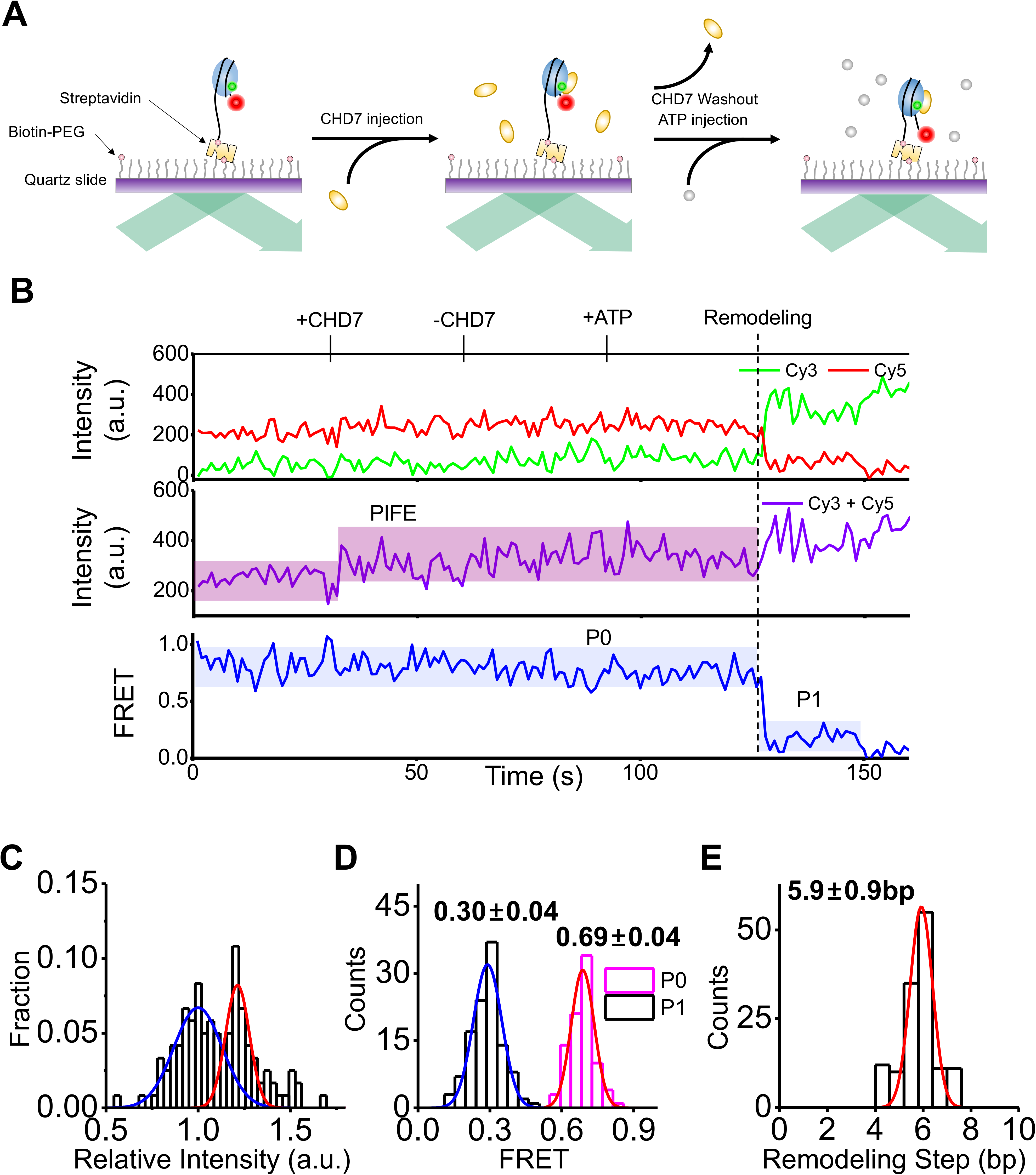
Single-molecule observation of nucleosome remodeling by CHD7. (A) Experimental scheme. Nucleosomes labeled with a FRET-pair (Cy3, green circle; Cy5, red circle) were immobilized on a microscope slide and CHD7 (200 nM) was injected into the chamber. After washing out free CHD7, ATP was injected into the reaction chamber to initiate chromatin remodeling. (B) Representative fluorescence intensity (top, green for Cy3 and red for Cy5), total intensity (middle, sum of Cy3 and Cy5), and FRET (bottom) time traces during nucleosome remodeling by CHD7. The trace shows total intensity increasing after CHD7 injection at 30 sec. CHD7 is washed out at 60 sec and ATP was added at 90 sec. This same color convention is used throughout the paper. (C) A histogram of total intensity(Figure 5B middle) between time 0 and 60 sec. The histogram is fitted to two Gaussian functions and x-axis is normalized by lesser intensity peak(blue). PIFE is distinguished by greater intensity peak (red). (D) FRET histograms of the initial state (P0) and the first translocation pause (P1) from 134 molecules which showed remodeling events. The histograms are fitted to Gaussian functions. (E) A histogram of remodeling step sizes and its fit to Gaussian function.

After establishing smFRET assays to monitor the binding and remodeling events of wild type CHD7, we studied the contribution of the N-CRD and the acidic pacth to the CHD7 activity using Nuc_APM_, CHD7_ΔN-CRD_ and CHD7_N-CRD_RA_ mutants. Interestingly, we observed two atypical traces in smFRET while the N-CRD CHD7 mutants remodel nucleosome, which were rarely observed with wild type CHD7. The remodeling events by the N-CRD deletions (CHD7_ΔN-CRD_) dominantly occurred with a clear slope (Fig. 6A) indicating that the deletion of the N-CRD substantially slows down the remodeling activity. Furthermore, the R to A mutations (CHD7_N-CRD_RA_) in the N-CRD lead to the another atypical FRET trace that the initial FRET drop was followed by FRET fluctuations between the high FRET and the low FRET states (Fig. 6B). These two atypical behaviors were not mutually exclusive and observed in the same molecules in some cases. However, the N-CRD deletion mutants showed more ‘Slow Remodeling’ while the point mutant showed more ‘Fluctuating’ FRET pattern. Fig. 6C shows how the portion of atypical behaviors affected by the CHD7 mutations. At this moment, it is not clear what is the exact nature of the remodeling event showing the fluctuating FRET pattern. However, it is conceivable that in the CHD7 mutants, ATPase activity is not efficiently coupled to proper nucleosome remodeling, causing a slow remodeling and non-fruitful remodeling exhibited by slope and fluctuating FRET pattern respectively. Overall these results strongly suggest that the N-CRD is required for the efficient remodeling activity.

**Figure 6.**
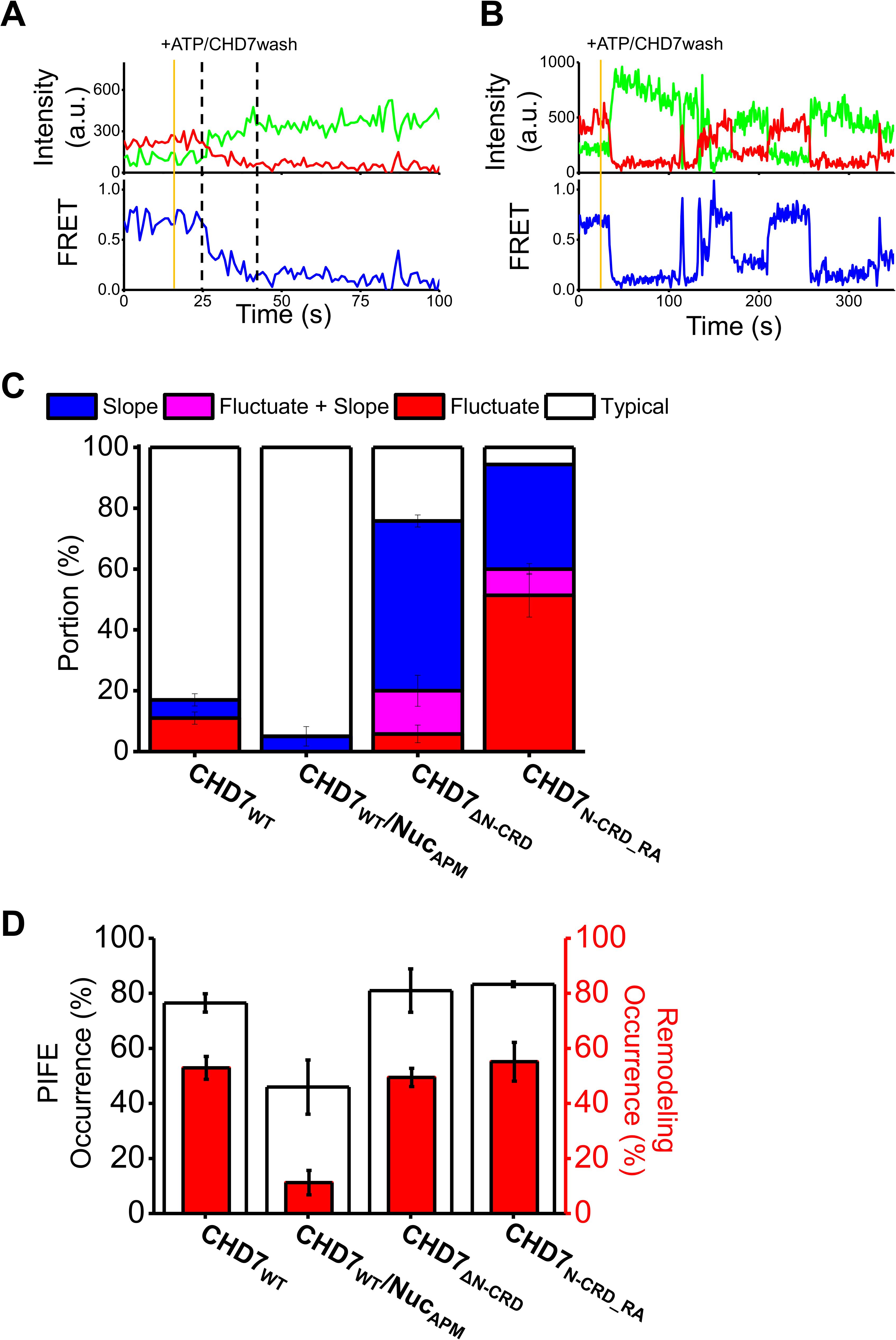
The mutational effects of the N-CRD domain of CHD7 and the acidic patch of nucleosome. (A) Representative fluorescent intensity and FRET time traces showing gradual remodeling of CHD7_ΔN-CRD_. Dashed lines indicate start and end of remodeling. Nucleosomes were incubated with CHD7_ΔN-CRD_ for 5 min before imaging. ATP is injected and free CHD7_ΔN-CRD_ is washed out at 15sec (orange). (B) Representative fluorescent intensity and FRET time traces showing remodeling fluctuations of CHD7_N-CRD_RA_. Nucleosomes were incubated with CHD7_N-CRD_RA_ for 5 min before imaging. ATP is injected and free CHD7_N-CRD_RA_ is washed out at 15sec (orange). (C) Portions molecule that exhibited remodeling events with a slope (blue), fluctuations (red), both a slope and fluctuations (magenta), and typical traces (white). CHD7_WT_/Nuc_APM_ indicates that APM nucleosome was used for the analysis. Otherwise, wild-type nucleosomes were used. The number of molecules used for the analysis were 82 for CHD7_WT_, 77 for CHD7_WT_/Nuc_APM_, 53 for CHD7_ΔN-CRD_, and 58 for CHD7_N-CRD_RA_. (D) Mutational effects on PIFE (white bar) and remodeling (red bar) occurrences. The ratios represent the portion of corresponding molecules among the molecules with the initial high FRET state.

When CHD7 is injected into the chamber where APM nucleosomes was immobilized, PIFE occurrence is substantially decreased compared with wild-type nucleosome (Fig. 6D). This observation suggests that the binding of CHD7 to nucleosome was compromised by the acidic patch mutations. Alternatively, CHD7 binds to APM nucleosomes in a way where PIFE does not occur due to the improper orientation of CHD7 to nucleosome. Furthermore, only 11.2% of the molecules among the molecules showing high FRET before ATP injection exhibited the remodeling activity compared to 52.9% of the wild type CHD7 (Fig. 6D). These data indicate that the CHD7 remodeling activity is substantially compromised by the acidic patch mutation despite CHD7 binds to APM nucleosome, indicating that the acidic patch of histones is important for CHD7 remodeling activity as well as nucleosome binding. Unexpectedly, the N-CRD CHD7 mutants showed comparable occurrence of PIFE to wild-type CHD7 indicating that the mutation/deletion in the N-CRD does not affect CHD7 binding to nucleosome, and they also show comparable remodeling occurence to wild-type when atypical remodeling traces were included (Fig. 6D). To further characterize the nucleosome remodeling by CHD7, we performed remodeling time analysis (Supplementary Fig. 10). Our analysis shows that the slow remodeling by CHD7 mutants is due to the slow kinetics of remodeling. Altogether, these results strongly suggest that the N-CRD is required for the efficient remodeling activity.

## Discussion

ATP chromatin remodelers form large multi-subunit complexes. CHD family is unique in that it is fully functional with only ATPase subunit (10,11). CHD7 has a Chromodomain followed by ATPase domain and a SANT DNA binding module (25). These domains are involved in the recognition of nucleosome substrates and their proper interaction with nucleosome renders the power-stroke by ATPase to properly remodel nucleosome (37,38). Here, we identify that arginine rich region in the N-terminal to Chromodomain likely interacts with the acidic patch of nucleosome and is critical for CHD7 activity. Our cryo-EM map of nucleosome complex with the N-CRD_Chromodomain shows the extra-density located above the acidic patch. Considering that the N-CRD alone without Chromodomain has comparable binding to the N-CRD_Chromodomain and Chromodomain alone does not bind to nucleosome (Fig. 1C), the extra-density observed is likely the N-CRD domain, suggesting that N-CRD domain interacts with the acidic patch. The binding of N-CRD domain to the acidic patch was further validated by the cryo-EM structure of N-CRD alone without a chromodomain bound to nucleosome with 40 bp linker DNA (Supplementary Fig. 5 and Supplementary Fig. 6). It should be noted that the sequence alignment of CHD7 orthologues among several species shows that the N-CRD arginine rich region is the only highly conserved region beside the Chromodomain at the N-terminal region of CHD7, suggesting the functional importance of the N-CRD.

The acidic patch is the major surface in nucleosome, which interacts with chromatin modifiers such as RCC1, Sir3 and DOT1L (27,28,31,34,35). The proteins recognizing the acidic patch contain arginine anchors composed of a stretch of arginine residues. The acidic patch interaction is shown to be critical for high-affinity binding, but also for proper orientation of proteins bound to nucleosome. For example, while mutating arginine anchor in Orc BAH domain compromises the interaction, mutations in the arginine anchor in DOT1L do not affect the binding *per-se* but affect the DOT1L activity (31).

Consistent with these previous findings, the CHD7 N-CRD domain involved in interacting the acidic patch contains a stretch of arginine, suggesting that CHD7 recognizes the acidic patch as other known acidic patch binding proteins. smFRET analysis shows that the mutations and deletion of the N-CRD does not affect the CHD7 binding in full-length context. This observation implies that the interaction between the N-CRD and nucleosome is likely critical for remodeling steps but not binding *per se* as in case of DOT1L (31). Interestingly, other ATP-dependent chromatin remodelers such as SWI/SNF and Swr1 complexes also interact with the acidic patch (32,33). However, these remodelers are composed of multiple subunits, and Ini1 and Arp6-Swc6 subunits in SWI/SNF and Swr1 complexes respectively are responsible for interacting with the acidic patch (33,39). In addition, a recent cryo-EM structure of SWI/SNF RSC complexes bound to nucleosome show that multiple-regions in the complex are involved in acidic patch binding. For example, in yeast SWI/SNF complex, SnAC region in the ATPases subunit (Sth1) and RPT region in Sfh1 are interacting with the acidic patch (40).

These observations suggest that CHD7 has evolved to contain all functional domain with in a single ATPase subunit rather to have multiple subunits. Interestingly, another CHD family remodeler, yeast Chd1p was shown that its remodeling activity is marginally affected by the acidic patch mutation (41). However, we observed that the acidic patch mutation greatly affects the CHD7 remodeling activity, suggesting that human CHD7 might function differently to yeast Chd1p.

Our smFRET analysis on the N-CRD mutants reveals its functional importance in the remodeling activity. For CHD7 remodeling activity, a power stroke by ATPase domain upon ATP hydrolysis is utilized to translocate DNA and CHD7 remodeler needs to hold histone octamer while translocating DNA in nucleosome. Therefore, it is imaginable that CHD7 utilizes the N-CRD domain to hold histone octamer via interacting with the acidic patch. In other ATPase remodelers, subunits in the complexes as mentioned above interact with the acidic patch and function as a ratchet for efficient remodeling. Consistent with this hypothesis, mutations or deletion in the N-CRD results in either slow or fluctuating remodeling activity. This arginine stretch in the N-CRD domain is found in other CHD remodeler belongting to subfamily III, but not subfamily I (Supplementary Fig. 11). In case of subfamily II, there is a double PHD domain at the N-terminal to Chromdomain. It would be interesting to see if the remodelers in the subfamily III would use a similar mechanism to remodel nucleosome. We also noticed that several mutations in the N-CRD domain, although not in the arginine stretch, are found in CHARGE syndrome, further emphasizing the functional importance of this domain (42–44).

In summary, this work identifies the arginine stretch located at the N-terminal to Chromodomain, which was not characterized previously, as a functionally important region interacting with nucleosome, and implies that chromatin remodelers are delicately regulated by several domains within ATPase subunit and/or subunits in the complexes.

## Materials and Methods

### Protein purification

CHD7 N-CRDs (590-940 and 800-940) were cloned into a modified pET28a vector having a TEV cleavage and expressed in *Escherichia coli* BL21(DE3) at 18 °C in the presence of 0.5 mM Isopropyl β-D-1-thiogalactopyranoside (IPTG) for 18 hrs. Cells were harvested in 300 mM NaCl, 50 mM Tris pH 8.0, 5 % Glycerol and 1 mM phenylmethylsulfonylfluoride (PMSF). The cells were lyzed and the lysates were cleared by centrifugation at 18,000 rpm (39,000 g) for 1 hr. His-tagged proteins were purified by Ni-NTA agarose affinity chromatography (Qiagen) and treated with TEV protease to remove the tag, followed by HiTrap Q HP (GE healthcare) and Superdex 200 26 600 (GE healthcare) in a buffer containing 150 mM NaCl, 50 mM Tris pH 8.0. Full-length CHD7 with N-terminal Flag tag and C-terminal His-tag was expressed and purified from Sf9 insect cell as described previously with minor modifications (25). Cells were harvested in a buffer containing 300 mM NaCl, 50 mM Tris pH 8.0 and 5 % Glycerol in presence of cOmplete™, Mini, EDTA-free Protease Inhibitor Cocktail (Roche) and lysed by 2 freeze-thaw cycles in liquid nitrogen. The lysate was then cleared by centrifugation at 18,000 rpm (39,000 g) for 90 min. The supernatant was incubated with M2-affinity resins (Sigma-aldrich) for 4 hrs and the proteins were eluted with 0.4 mg/ml FLAG peptide. The eluted samples were applied to Ni-NTA resins in presence of 5 mM Imidazole. After washing the resins with a buffer containing 50 mM Tris pH 8.0 and 500 mM NaCl followed by 300 mM NaCl buffer, full length CHD7 was eluted with 150 mM NaCl, 20 mM HEPES pH 7.5 and 200 mM Imidazole. The protein was concentrated using Amicon Ultra centrifugal filter devices (Millipore). CHD7_ΔN-CRD_ and CHD7_N-CRD_RA_ were also purified as the wild type full length CHD7.

### Limited proteolysis

3 μg of proteins in 150 mM NaCl and 50 mM Tris-HCl (pH 8.0) were incubated with increasing amount of trypsin at 27 °C for 30 mins. The reactions were stopped by the addition of SDS-PAGE sample loading buffer. The samples were analyzed by 4-20 % gradient SDS-PAGE gel (Bio-Rad) stained with coomassie blue.

### EMSA assay

5 μM of nucleosomes were mixed with increasing amounts of CHD7 enzymes (concentration specified in each figure) in 50 mM NaCl and 20 mM HEPES pH 7.5. In case of free DNA fragments, 1 μM of DNA was used. All the mixtures were incubated on the ice for 1 hr and analysed with 4 % native PAGE (0.5 × TBE) followed by staining with Red safe (Intron). The amount of bound nucleosomes and DNA was measured with ChemiDoc MP (Bio-rad) using Image Lab software.

### REA assay

REA assays were performed as described previously with minor modification (45). The MfeI (NEB) site located at 28 bp in the nucleosome was used to measure the remodeling activity. REA reactions were performed at 30 °C in REA buffer (12 mM HEPES pH 7.5, 4 mM Tris pH 7.5, 50 mM KCl, 10 mM MgCl_2_, 10 % glycerol, 0.02 % NP-40). 50 nM of CHD7 was incubated with 3 U/ʎ of MfeI for 7 min and after adding the ATP was incubated for 2 more minutes. In case of assay using CHD7_N-CRD_RA_, 1.5 U/ʎ of MfeI was used. The reaction was initiated by adding 20 nM of nucleosomes. Time points were quenched with 1.5 X volume of quenching buffer (10 % Glycerol, 70 mM EDTA pH 8.0, 20 mM Tris pH 7.5, 2 % SDS, a pinch of bromophenol blue). Samples were deproteinized with 0.5 mg/ml of Protease K (NEB) and incubated for 1 hrs at 37 °C. Cut DNA was separated from uncut DNA on 8 % native PAGE (0.5 × TBE) followed by Red safe (Intron) staining

### ATPase assays

The EnzChek Phosphate assay kit (Thermo Fisher Scientific) was used to measure steady state ATPase rates of CHD7 and mutants. 100 nM of CHD7 was incubated in assay buffer (50 mM NaCl, 50 mM Tris pH 8.0) with or without saturating concentration of substrate (300 nM : nucleosome, nucleosome+80 and DNA fragment) and 1 mM MgATP was added to initiate the reaction at 30 °C. Absorbance at 360 nm was monitored on a Spark multimode microplate reader (Tecan) 5 sec after initiating the reaction, and with 10 sec intervals thereafter for 40 min.

### Chemical crosslinking and MS analysis

Purified full-length CHD7 and nucleosome with 40 bp of linker DNA were mixed with molar ratio 3:1 in presence of 1 mM MgCl_2_ and 1 mM ADP.BeFx. Final 1 mM of disuccinimidyl suberate (DSS)-H12/D12 (Creative molecules) was added and incubated the mixture for 30 min at 37 °C with mild shaking. The reaction was quenched by adding final 50 mM of ammonium bicarbonate incubating for 20 min at 37 °C. The samples were denatured with 8 M urea in the presence of 2.5 mM TCEP. The free cysteine thiols were blocked by adding final 5 mM of Iodoacetate. The samples were digested with Lysyl Endopeptidase incubating at 37 °C for 5 hrs at 1:100 enzyme to substrate ratio and followed by second digestion with Trypsin for overnight at 1:50 enzyme to substrate ratio. The reaction was stopped by adding final 2 % (v/v) of formic acid. This peptides were cleaned with Sep-Pak column in water/Acetonitrile/Formic acid =50:50:0.1. The cross linked peptides were enriched by Superdex peptide PC 3.2/30 in water/Acetonitrile/Trifluoroacetic acid=70:30:0.1 as mobile phase. Fractions were collected, dried and analysed at Taplin Mass Spectrometry Facility at Harvard Medical School. Data analysis was performed by xQuest (46) and an xQuest LD score cutoff of 12 was selected.

### Cryo-electron microscopy

#### Sample preparation

Nucleosome core (Nuc) was prepared with Xenopus histones with 601 147 bp DNA as previously described (31). Nucleosome (Nuc+40) with 40 bp extra DNA was prepared similary using 601+40 187 bp DNA. CHD7_N-CRD-CD_ (590-940 a.a.) and CHD7_N-CRD_ were mixed with Nuc and Nuc+40 in 5:1 ratio in 50 mM NaCl and 20 mM HEPES pH 7.5 respectivly. The samples were cross linked with 0.1 % Glutaraldehyde by incubating on ice for 10 min. The reactions were quenched by adding 5 mM Lysine and 100 mM Tris pH 7.5. To purify the homogeneous population, sucrose gradient separation was performed by ultra-centrifugation at 30,000 rpm (85,000 g), 16 hr using a Beckman SW55 rotor in a condition of 5-20 % sucrose gradient. A 3 μl drop of samples were applied to glow-discharged QuantiFoil™ R2/2 200 mesh. After 20 sec incubation, grids were blotted for 4 sec, and then plunge-frozen in liquid ethane using a Vitrobot 4 (Thermo Fisher).

#### Data collection and analysis

For CHD7_N-CRD-CD_ Nuc complex structure, micrographs were collected at SciLifeLab in Stockholm on Titan krios (Thermo Fisher) operated at 300 KeV, equipped with K2 summit direct detector (Gatan). In total, 9,369 micrographs were recorded using −1.2~2.2 μm defocus range at pixel size 0.82 Å /pixels and with a total electron dose of 47.94 e^−^/Å^2^. Details are summarized in Supplementary Table 2. Motion correction was carried out during data collection using MotionCor2 (47) and CTF parameters were estimated using CTFFIND4 (48). Image processing was performed with Relion 3.0 (26). Details of data processing schemes are shown in Supplementary Fig. 3. Total 1,859,258 particles were picked using automated particle picking and extracted with box size of 230 pixels. Bad particles were removed by serial 2D classification, yielding a total 946,655 particles. First round of 3D classification was performed using use fast subset option with a model of free nucleosome generated from x-ray crystallography structure (PDB ID 1KX5) as reference. Well aligned classes were selected and total 328,395 particles were obtained. This particles were further subjected to additional rounds of 3D classification using the newly generated model in each round as a reference. Final 17,743 particles were used for 3D refinement and post processing with automatic B-factor. The refined cryo-EM structure showed 7.25 Å resolution as determined by the gold standard FSC at 0.143 criterion.

For the CHD7_N-CRD_ and Nuc+40 complex structure, micrographs were collected at KAIST Center for Research Advancement (KARA), Daejeon, Korea using a Glacios (Thermo Fisher) operated at 200 KeV, equipped with a Falcon III direct detector. In total, 1,782 micrographs were recorded using - 1.2~3.2 μm defocus range at pixel size 1.144 Å /pixels and with a total electron dose of 39.54 e^−^/Å^2^. Details are summarized in Supplementary Table 2. All the image processing was performed using RELION-3.0 (26). Details of data processing schemes are shown Supplementary Fig. 5. The movie stack was aligned with MotionCorr2 (47), and CTF was corrected with CTFFIND4 (48). Total 484,194 particles were picked using automated particle picking and subjected to 2D class averaging. Poorly aligned particles were removed by serial 2D classification followed by three rounds of 3D classification. Final 47,550 particles were used for 3D refinement and post processing with automatic B-factor. The refined cryo-EM structure showed 7.14 Å resolution as determined by the gold standard FSC at 0.143 criterion.

### sm FRET

The standard nucleosomes used in single molecule FRET experiment were designed to have a 78-bp spacer on the entry side and a 3-bp spacer on the exit side (Supplementary Fig. 5A). DNA fragments containing 601 nucleosome positioning sequence(49) were generated by PCR as described (50). Primers with Cy5 at the 5’ end or biotin were purchased from IDT. PCR products were purified by 5 % native PAGE gel. Nucleosomes were reconstituted with histone octamer containing either Cy3-labeled H2A T120C and Cy5-labeled DNA fragment using salt gradient dialysis. Quartz slides and coverslips were cleaned with piranha solution (mixture of 3:1 concentrated sulfuric acid:30%[v/v] hydrogen peroxide solution), coated with aminopropyl silane first, and then with mixture of PEG (m-PEG-5000, Laysan Bio) and biotin-PEG (biotin-PEG-5000, Laysan Bio). A flow cell was assembled by combining a coverslip and a quartz slide with double-sided adhesive tape (3 M). For convenient buffer exchange, polyethylene tubes (PE50; Becton Dickinson) were connected to the flow cell. Nucleosome substrates were immobilized on a PEGylated surface using biotin-streptavidin conjugation. smFRET experiments were performed in an imaging buffer (10 mM Tris-HCl pH 8.0, 100 mM KCl, 3 mM MgCl_2_, 5 mM NaCl) containing a gloxy oxygen scavenging system (a mixture of glucose oxidase (1 mg/ml, Sigma), catalase (0.04 mg/ml, Sigma), glucose (0.4 % (w/v), Sigma), and Trolox (2 mM, Sigma)). Single-molecule fluorescence images were acquired at a frame rate of 10 Hz using a home-built prism-type total internal reflection fluorescence microscope equipped with an electron-multiplying charge coupled device camera (Ixon DV897; Andor Technology). Cy3 and Cy5 were alternately excited with 532 nm and 633 nm lasers using the ALEX(Alternative Laser Excitation) technique (51). Experimental temperature was maintained at 30 °C using a temperature control system (Live Cell Instruments). Data acquisition and FRET trace extraction were done using homemade programs written in LabView (National Instruments) and IDL (ITT), respectively. The intensity traces of Cy3 and Cy5 were analyzed using a custom made Matlab(MathWorks) script. The FRET efficiencies were calculated from the donor (ID) and acceptor (IA) fluorescence intensities as EFRET = IA/(IA + ID).

## Data availability

The cryo-EM maps of N-CRD_Chromdomain core nucleosome and N-CRD nucleosome+40 were deposited at the EM database as EMD-30298 and EMD-30630 respectively.

## Acknowledgements

We thank the members of the Song Lab for helpful discussions. We also thank the staff at the SciLifeLab and KARA for the Data collection. We thank Alexander Leitner at ETH Zurich for helpful discussion regarding XL-MS. The data processing was performed at Global Science experimental Data hub Center (GSDC) at Korea Institute of Science and Technology Information (KISTI). This work is partially supported by grants (NRF-2020R1A2B5B03001517, NRF-2016K1A1A2912057 to J.S. and NRF-2019R1A2C2005209 to S.H.) from the National Research Foundation of Korea, and a grant from the Swedish Research Council (VR-2016-03810 to H.H. and P.H.).

## Author contributions

E.L., C.K., K.B., S.H., J.S. conceived the idea. E.L. performed the binding and remodeling analysis, and E.L., P.H., H.H. performed cryo-EM. C.K. S.H. performed single-molecule FRET analysis. All authors examined the data and wrote the manuscript.

---

**Supplementary Figure 1.**
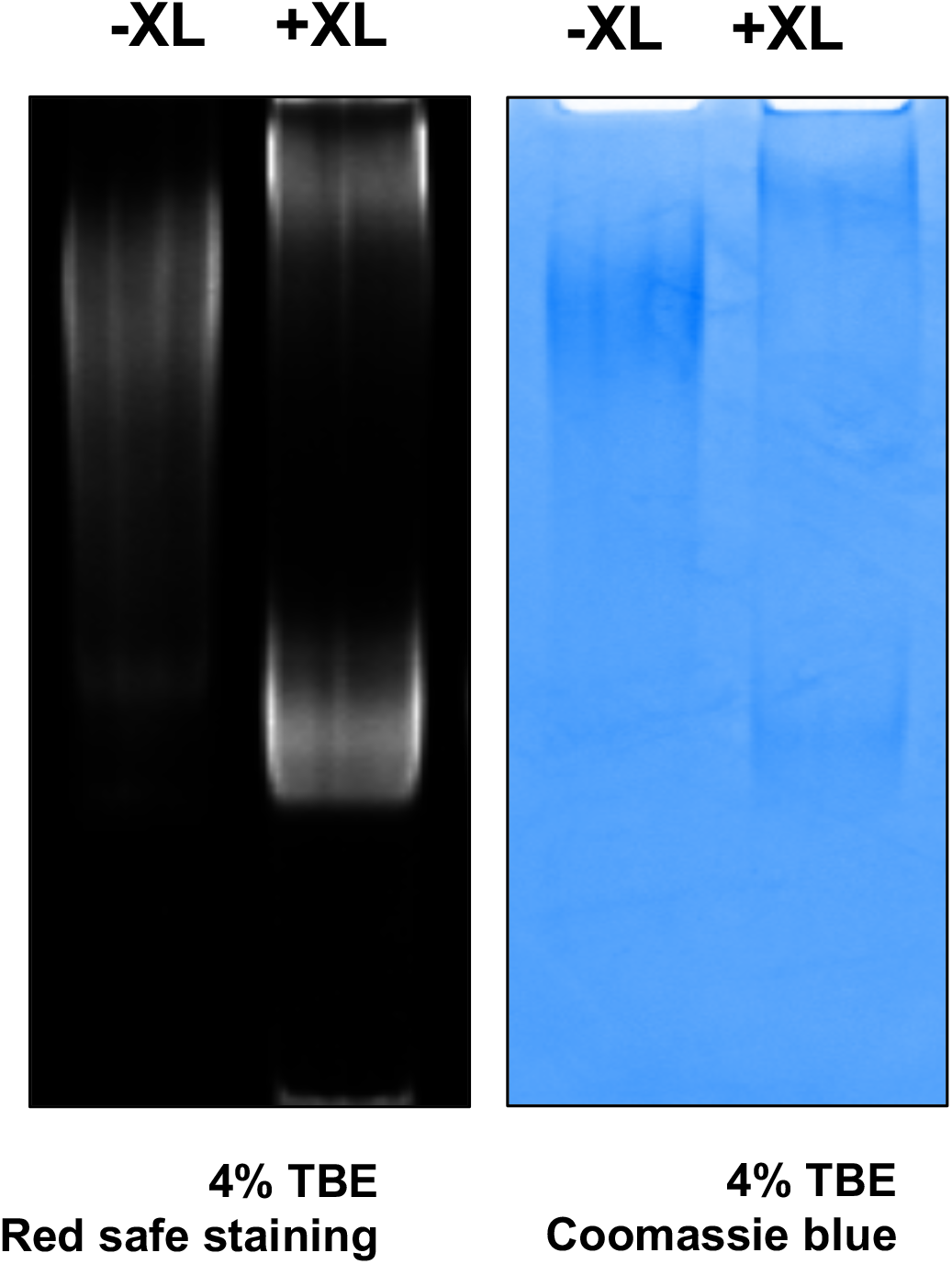
A sample preparation for cross-linking mass spectrometry. The quality of DSS cross-linked CHD7 nucleosome complex was analyzed with 4% native PAGE showing the homogeneity of the sample. Un-crosslinked and crosslinked samples were indicated with −XL and +XL respectively.

**Supplementary Figure 2.**
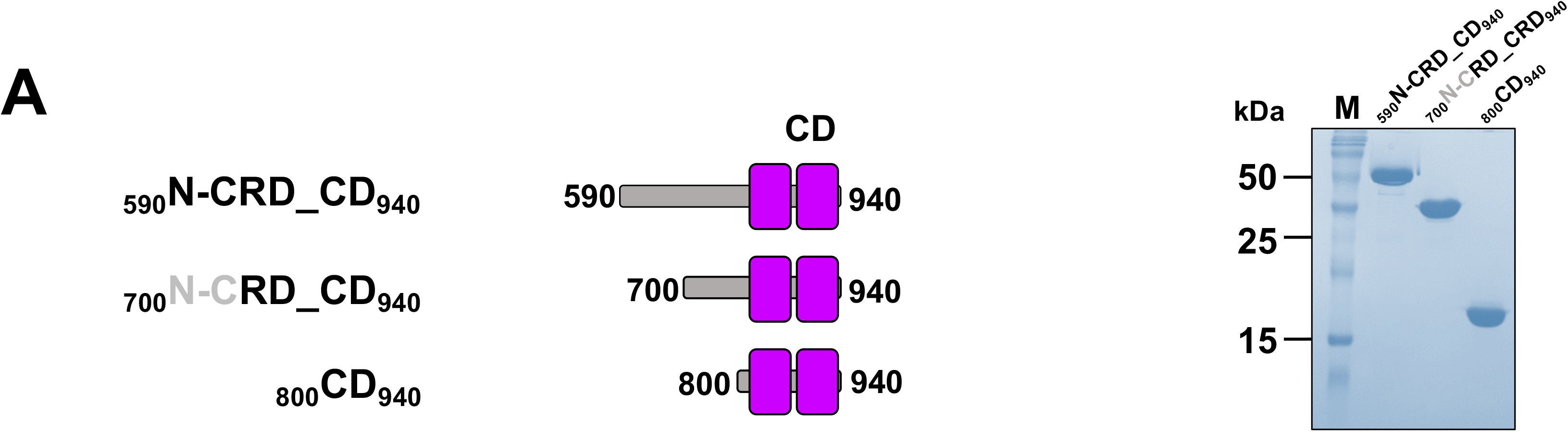
EMSA analysis of N-CRD_Chromodomain binding to substrates. (A) The quality of the proteins used in gel shift assay was analyzed by SDS-PAGE with coomassie blue staining. CD stands for Chromodomiain.

**Supplementary Figure 3.**
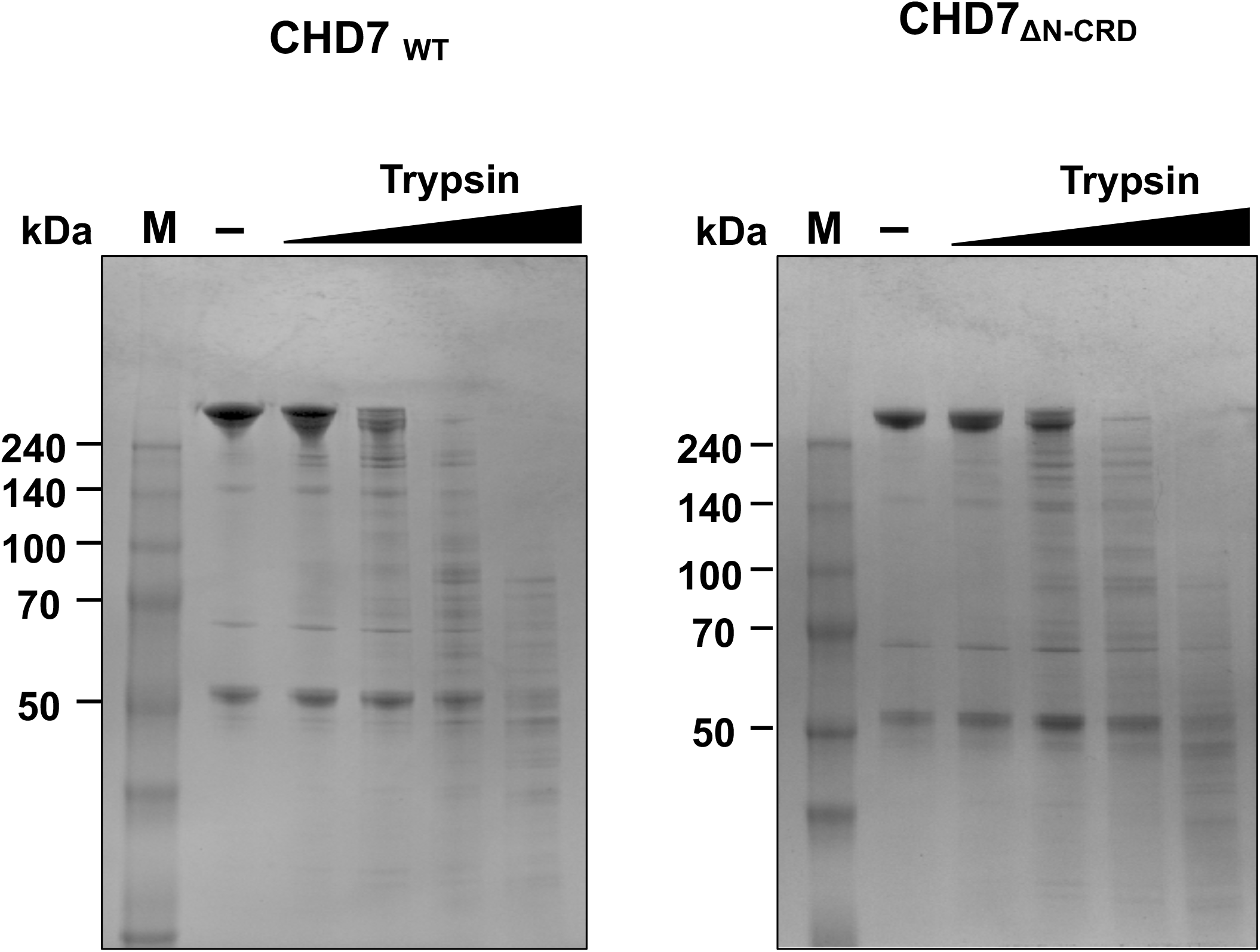
Limited proteolysis analysis of CHD7_WT_ and CHD7_ΔN-CRD_. CHD7_WT_ and CHD7_ΔN-CRD_ were each incubated with the increasing amounts of trypsin and the reactions were analyzed by SDS-PAGE gel followed by a coomassie blue staining. The limited proteolysis analysis shows the similar cleavage pattern between CHD7_WT_ and CHD7_ΔN-CRD_, indicating that the deletion of N-CRD does not causes misfolding of the protein.

**Supplementary Figure 4.**
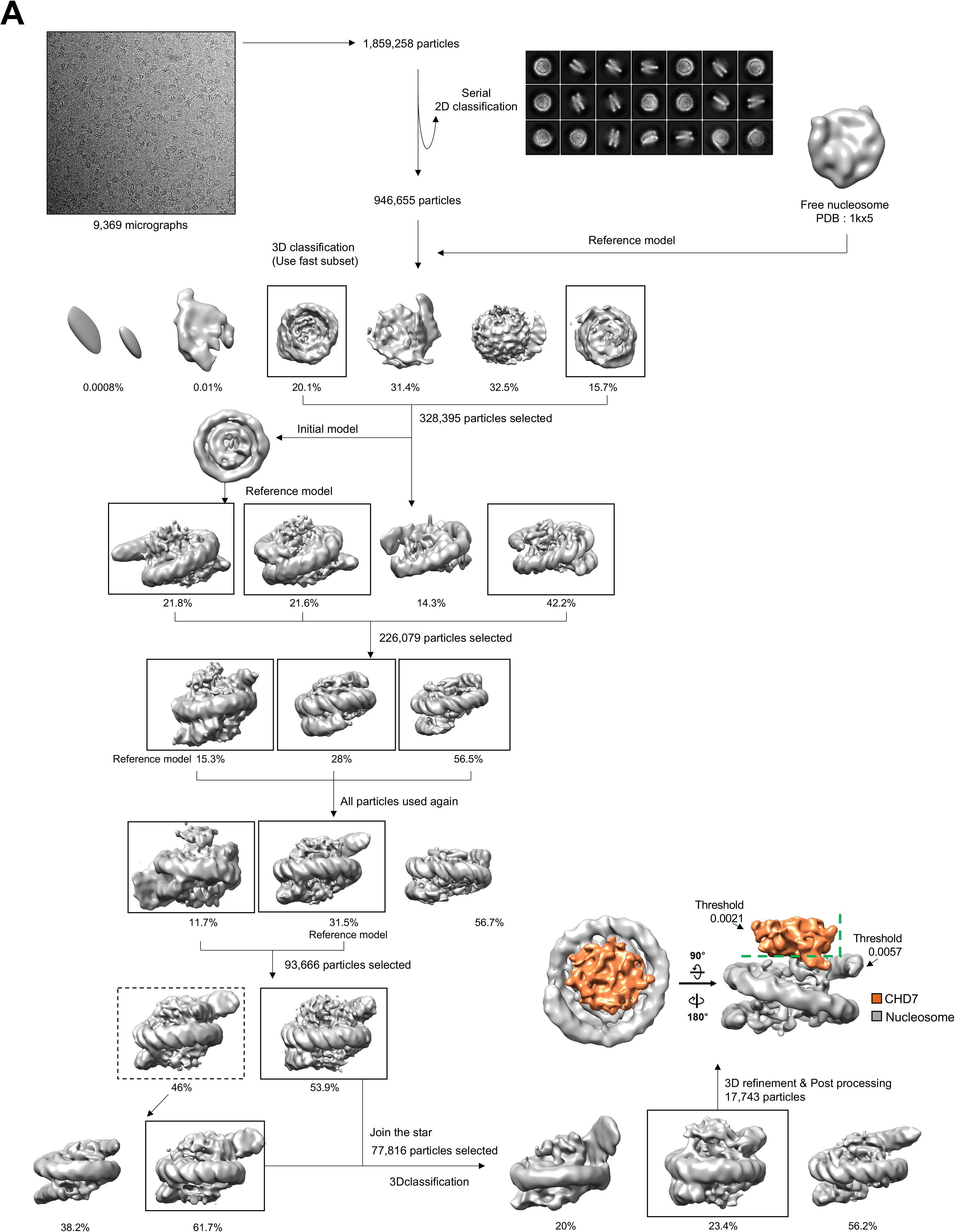
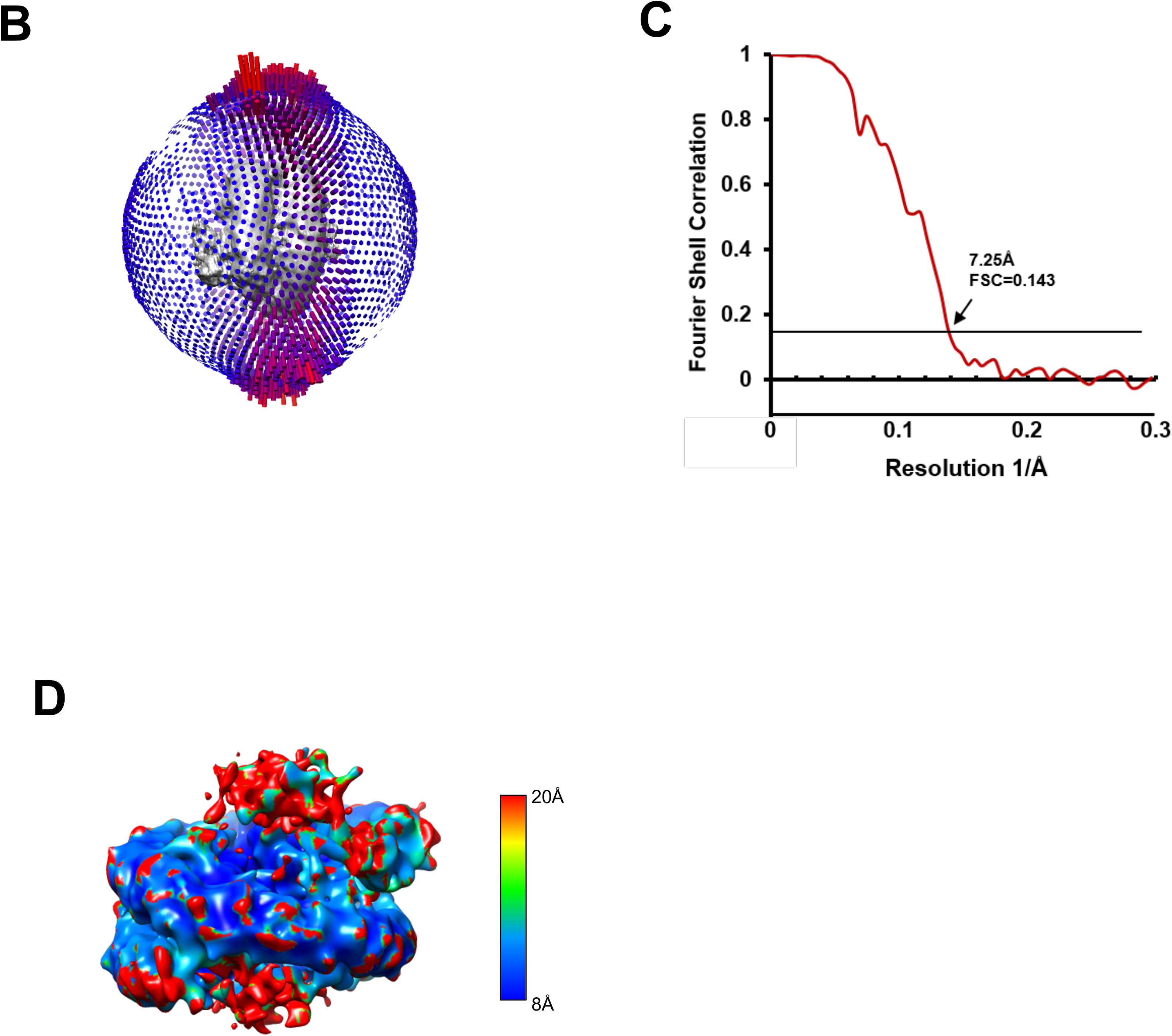
Cryo-EM processing of N-CRD_Chromo nucleosome core complex. (A) A detailed scheme for the cryo-EM data processing of N-CRD_Chromo Nucleosome complex. All processing was performed with Relion3.0 (B) An Euler angle distribution of the particles used for the reconstruction (C) The FSC curve indicates that 7.25 Å resolution by the gold standard FSC at 0.143 criterion (D) A local resolution on the map by ResMap

**Supplementary Figure 5.**
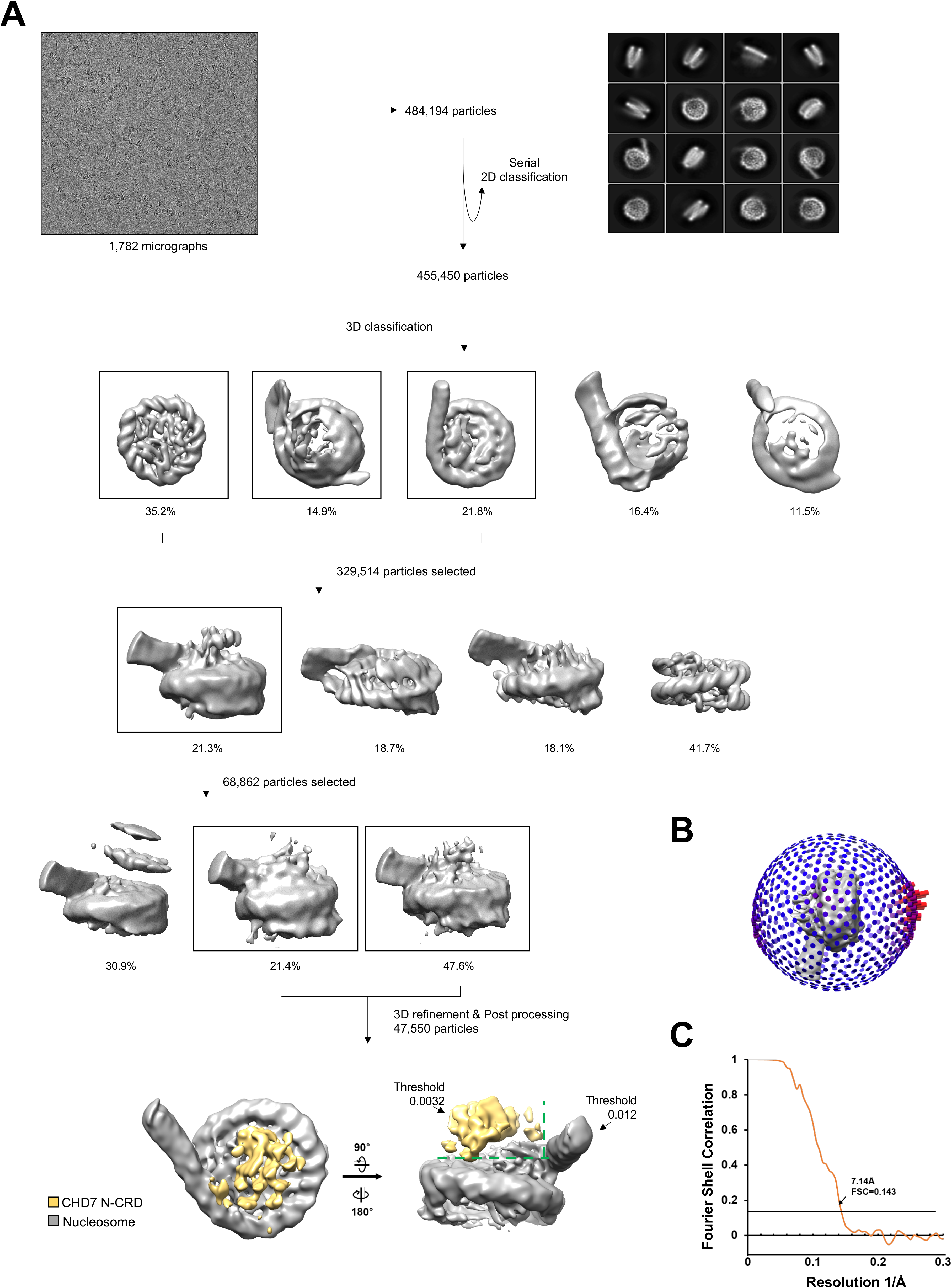
Cryo-EM processing of CHD7 N-CRD and Nucleosome with 66 extra linker DNA complex. (A) A detailed scheme for the cryo-EM data processing of N-CRD Nucleosome complex. All processing was performed with Relion3.0 (B) An Euler angle distribution of the particles used for the reconstruction (C) The FSC curve indicates that 7.14 Å resolution by the gold standard FSC at 0.143 criterion

**Supplementary Figure 6.**
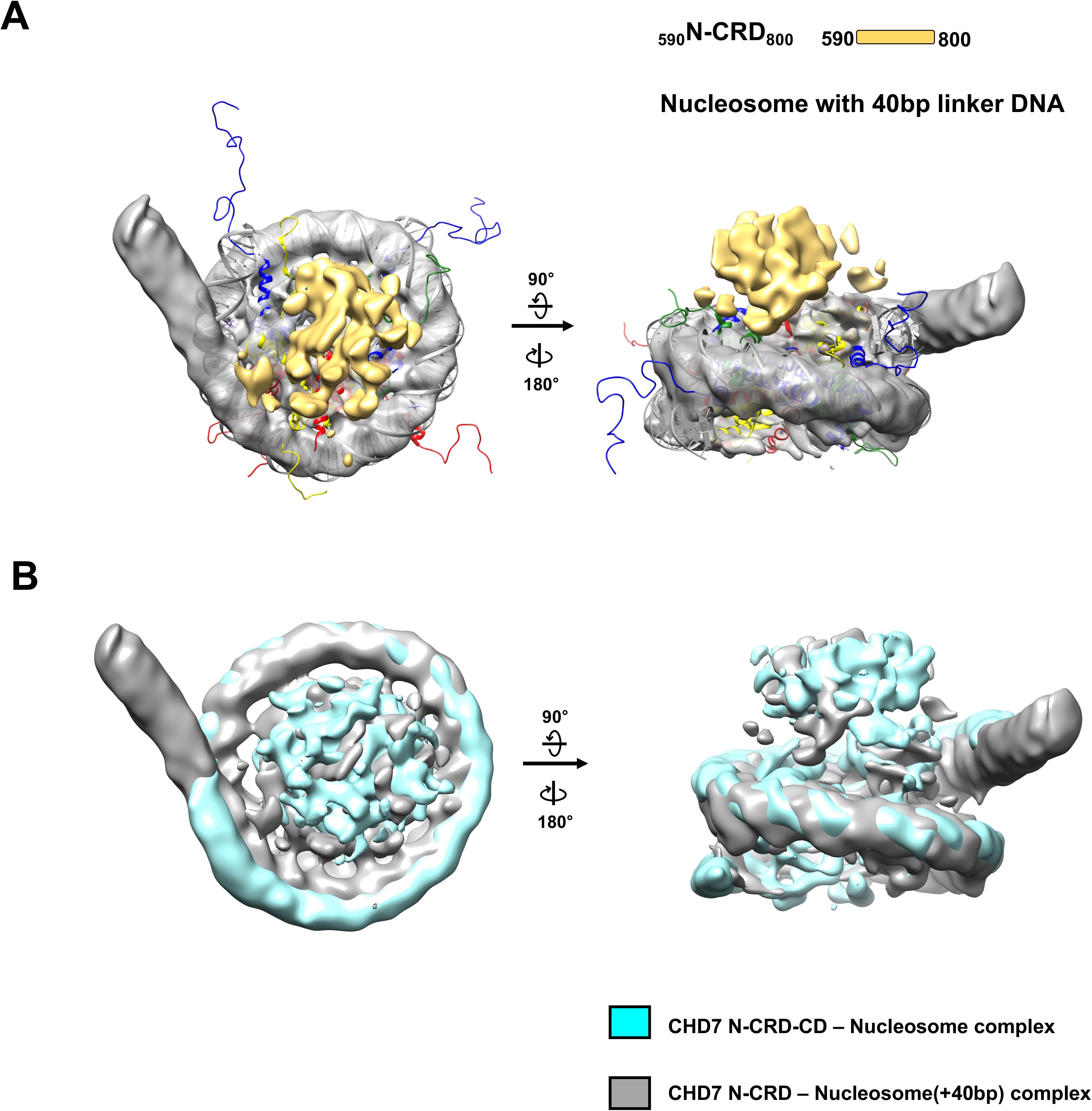
The Cryo-EM map of N-CRD bound to nucleosome with 40bp linker DNA. (A) The Cryo-EM map for nucleosome is shown in gray and the crystal structure of nucleosome (1KX5) was docked in the map. The extra-density for N-CRD is shown in yellow. The map was drawn at a contour 0.012 for nucleosome and 0.0032 for N-CRD respectively. (A) A superimposition of the cryo-EM maps between N-CRD-CD – Nuc core complex and N-CRD Nuc(+40) complex. The Cryo-EM map for N-CRD-CD – Nuc complex is shown in Cyan and N-CRD – Nuc(+40) complex is shown in gray.

**Supplementary Figure 7.**
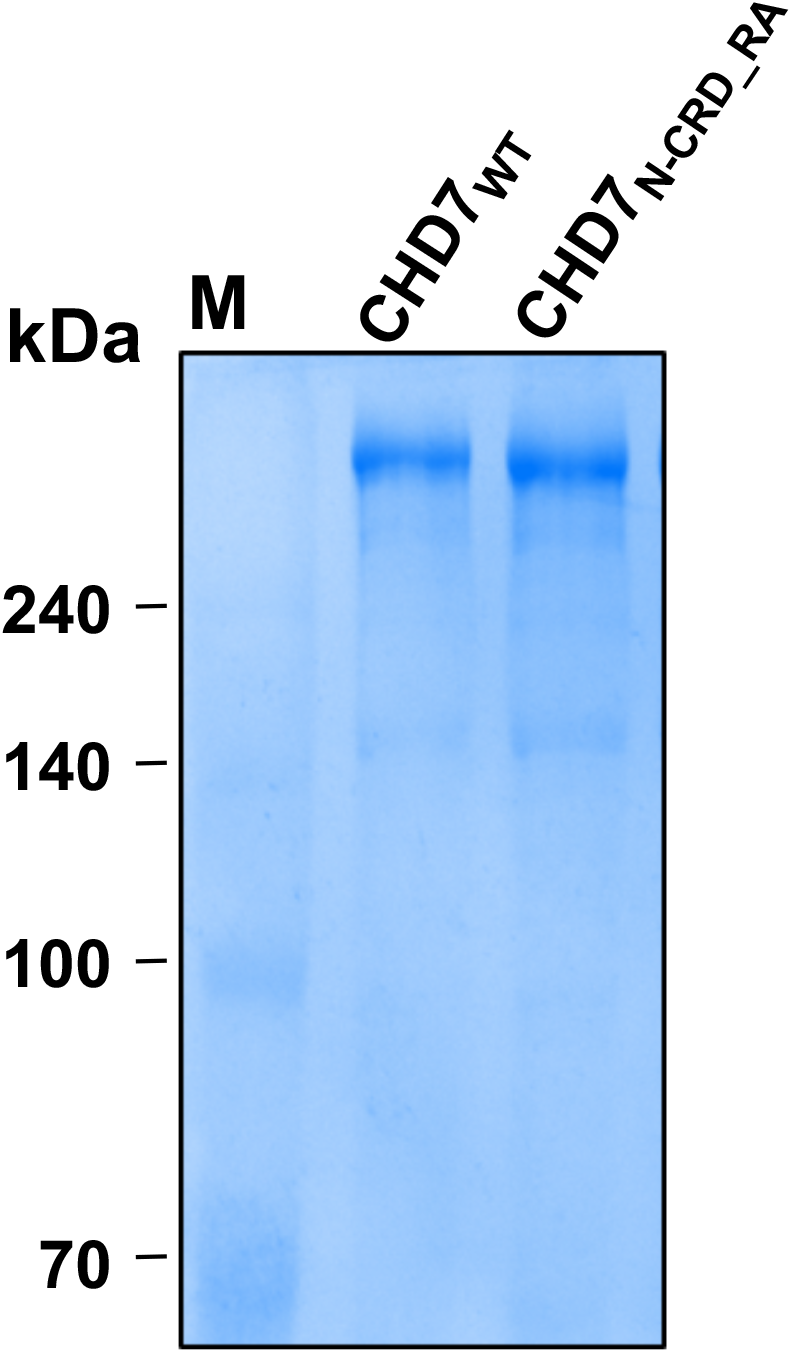
SDS-PAGE analysis of CHD7 proteins. The qualities of the proteins were analyzed by SDS-PAGE stained with coomassie blue

**Supplementary Figure 8.**
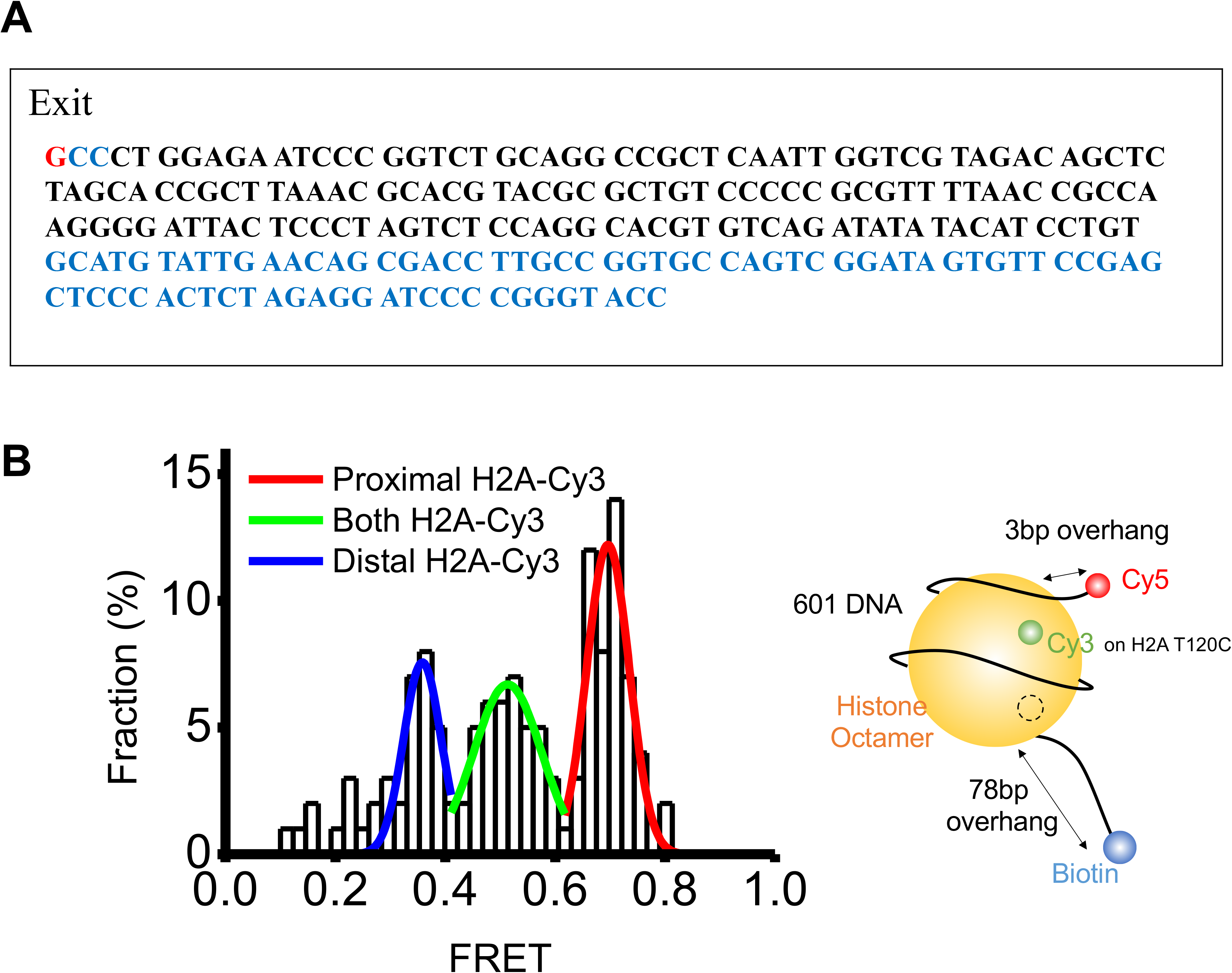
DNA design and molecule selection scheme for smFRET. (A) DNA sequences used in the study. (A) The DNA fragment consists of 601 nucleosome positioning sequence (black) flanked with the 3 bp linker on the exit side and the 78 bp on the entry side (blue). The Cy5 labeling position is marked in red. (B) FRET histogram obtained from surface-immobilized nucleosomes (left, n = 200) and a schematic representation of the nucleosome species corresponding to the high FRET state (right). The histogram shows three FRET peaks. The low and high FRET correspond to species labeled at the distal and proximal H2A, respectively. The middle FRET corresponds to doubly-labeled species. We selected only high FRET species for data analysis.

**Supplementary Figure 9.**
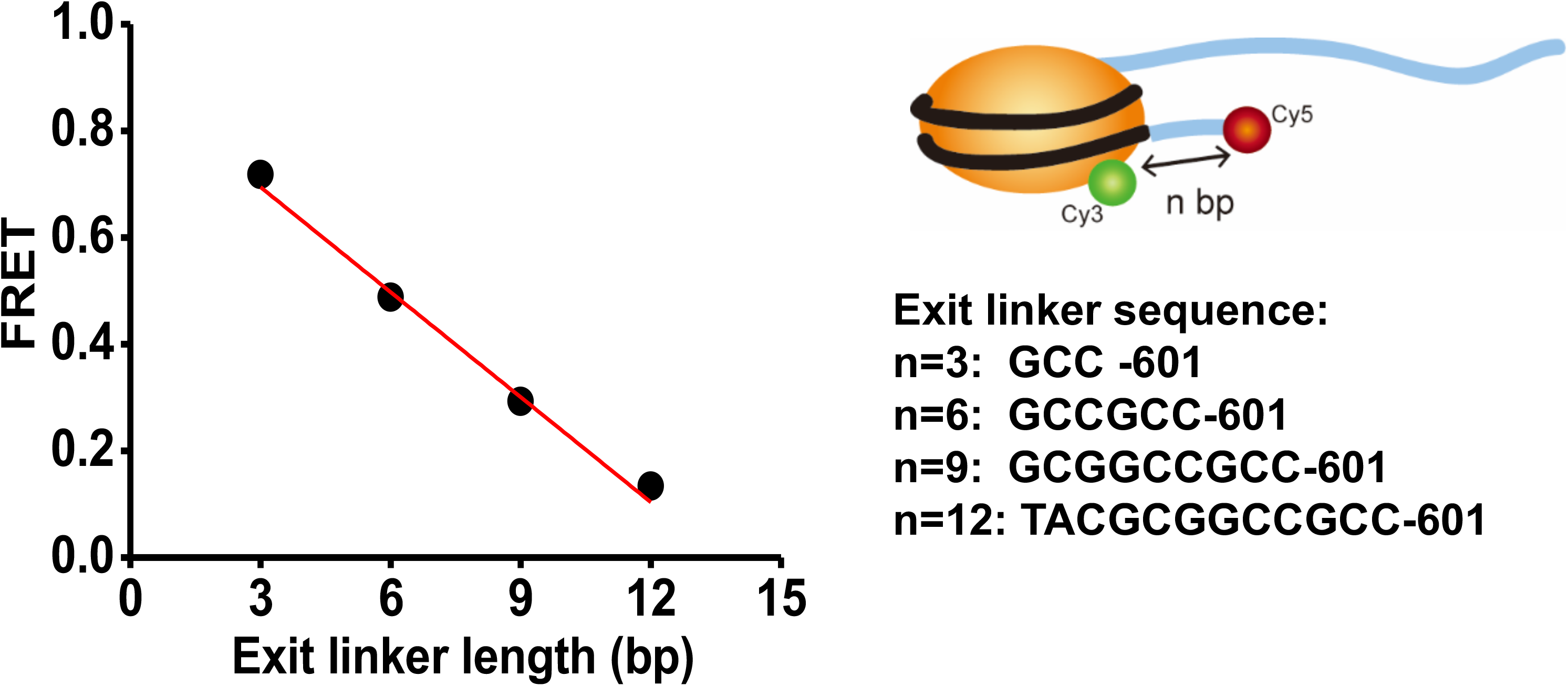
Calibration data for step size determination. FRET was obtained from nucleosomes with varying linker DNA lengths (n= 3, 6, 9, 12 bps) on the exit side. The linear-fit of the data had a slope of −0.066 ± 0.0004 per base pair.

**Supplementary Figure 10.**
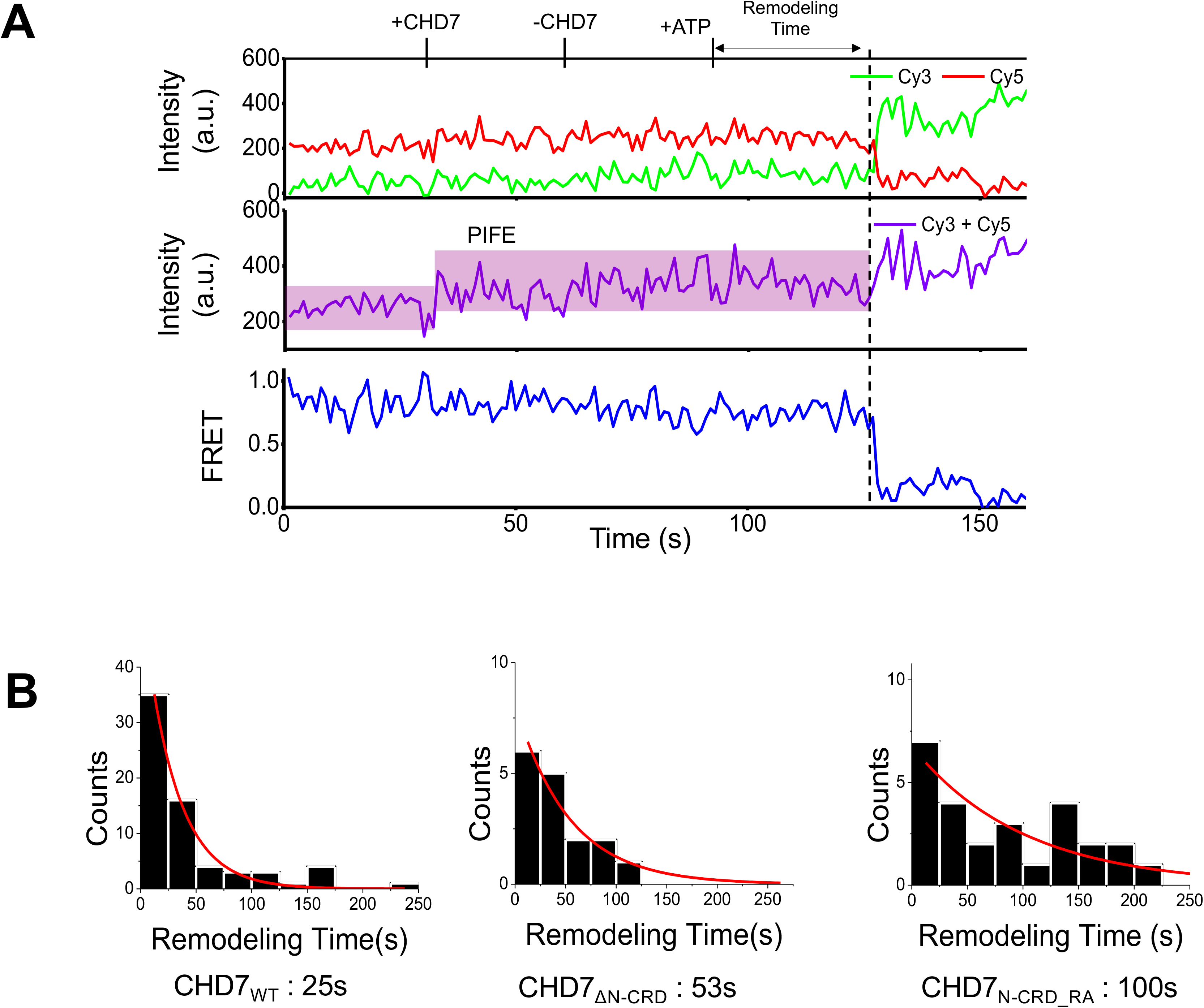
Remodeling time analysis. (A) A kinetic parameter; remodeling time, the delay between ATP injection and the remodeling. (B) Histograms of remodeling time of various CHD7 WT and mutants. The histogram is fitted to single exponential decay and t_0_ are described below. The number of molecules used for the analysis were 67 for CHD7_WT_,16 for CHD7_ΔN-CRD_ and 26 for CHD7_N-CRD_RA_.

**Supplementary Figure 11.**
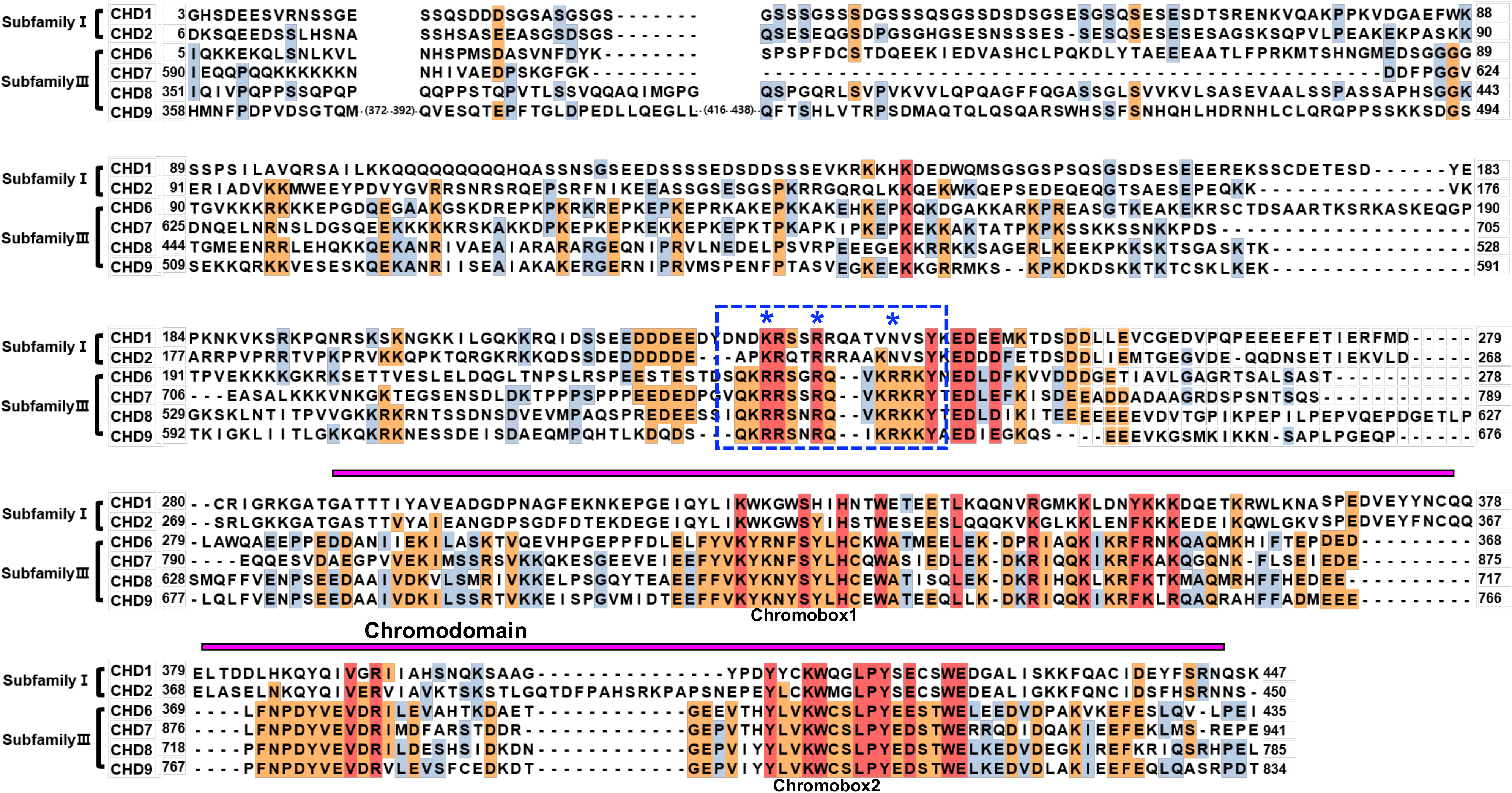
A sequence alignment of N-CRD_Chromodomain among CHD subfamily I and III. A sequence alignment of N-CRD_Chromodomain among other CHD remodelers belonging to subfamilyⅠand subfamilyⅢ. In case of subfamilyⅡ, since the PHD domain exists in front of the chromodomain, it was excluded from alignment. A blue dotted box shows the arginine rich stretch in N-CRD. The mutated arginines are marked with asterisks.

**Supplementary Table 1.**
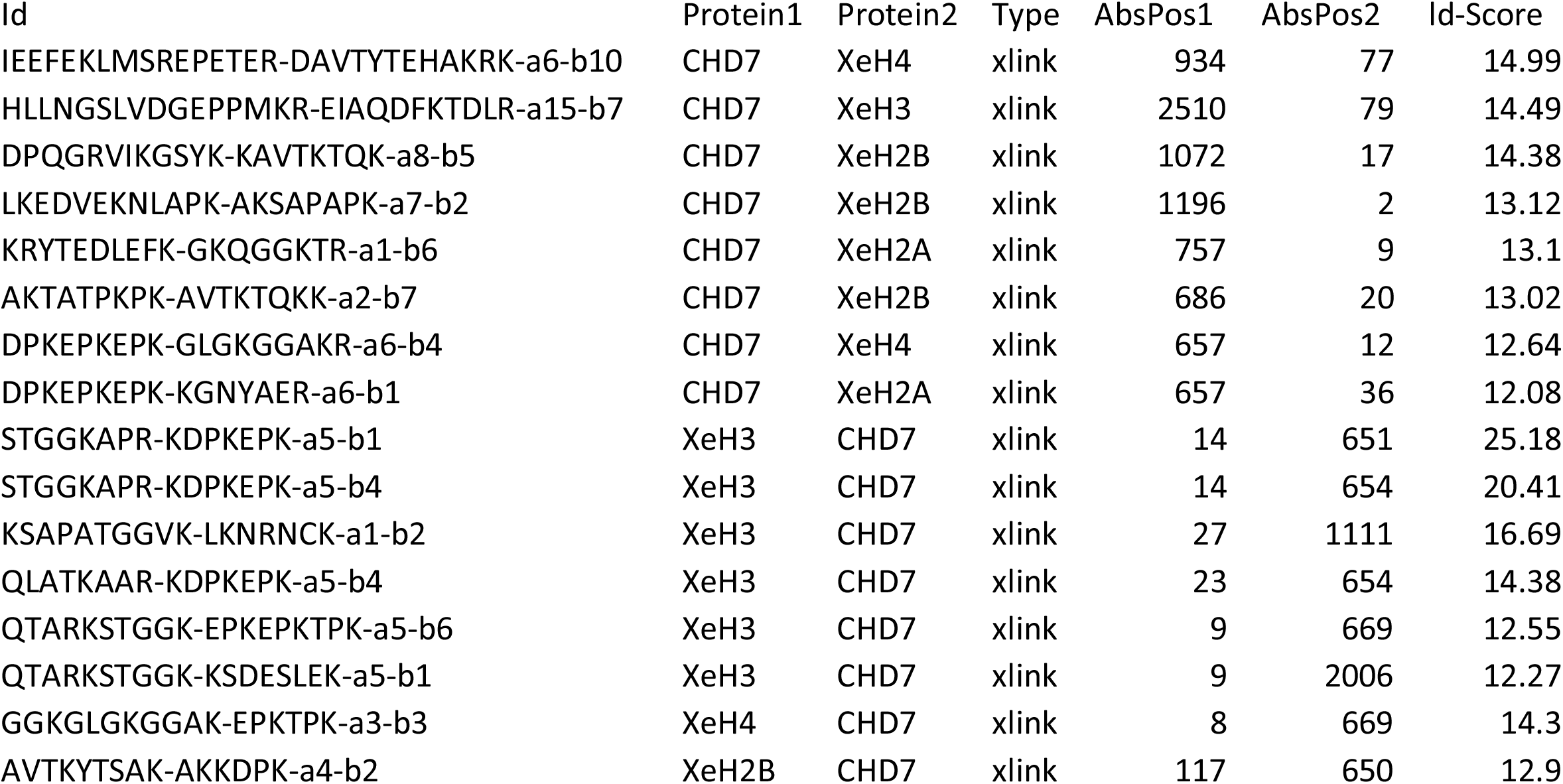
A list of crosslinks from XL-MS.

**Supplementary Table 2.**
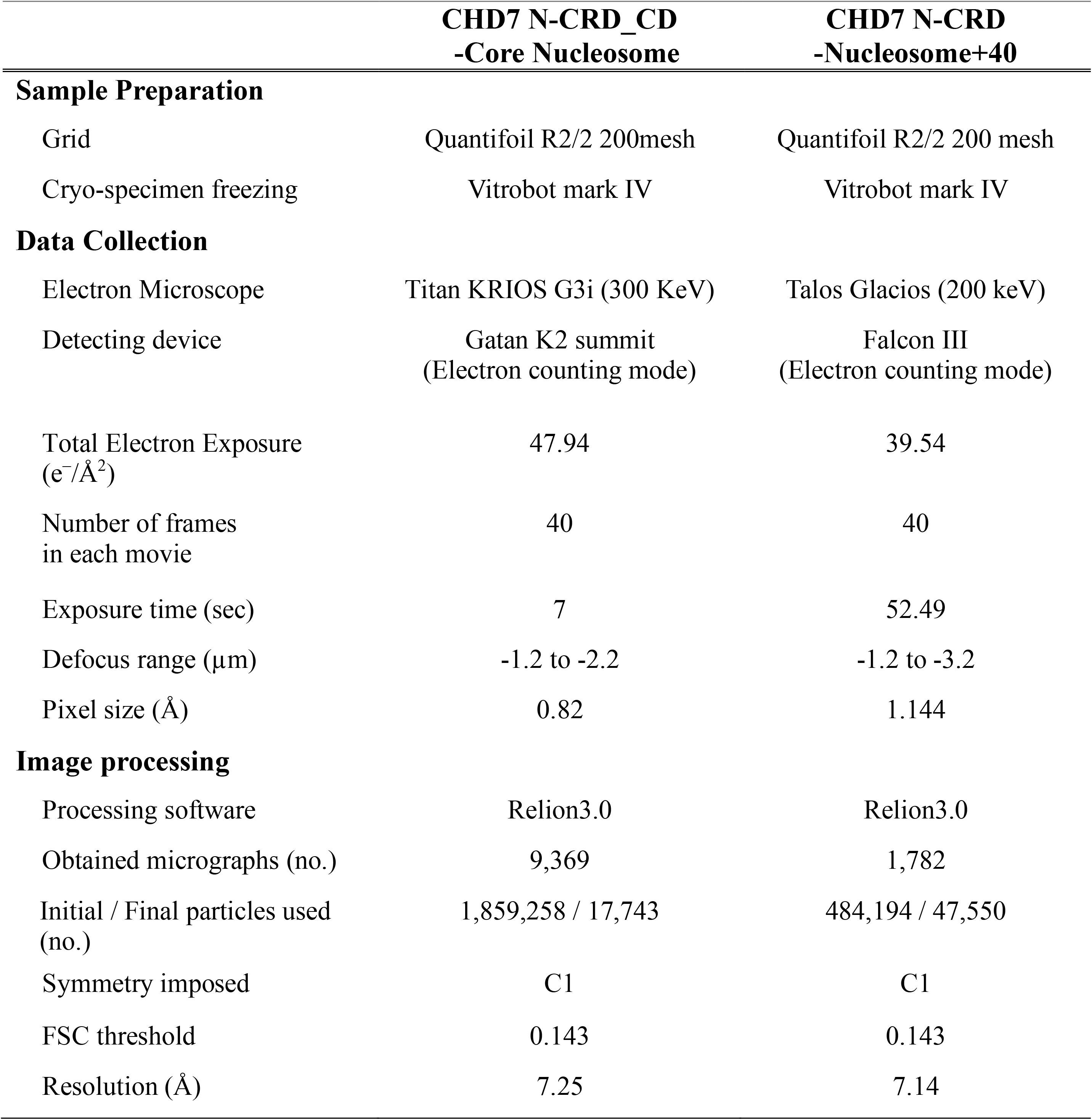
Cryo-EM data collection and statistics.

